# Hyperphosphorylated tau Inflicts Intracellular Stress Responses That Are Mitigated by Apomorphine

**DOI:** 10.1101/2023.05.13.540661

**Authors:** Zhenfeng Song, Kuang-Wei Wang, Hsiao-Tien Chien Hagar, Hong-Ru Chen, Chia-Yi Kuan, Kezhong Zhang, Min-Hao Kuo

## Abstract

**Background:** Abnormal phosphorylation of the microtubule-binding protein tau in the brain is a key pathological marker for Alzheimer’s disease and additional neurodegenerative tauopathies. However, how hyperphosphorylated tau causes cellular dysfunction or death that underlie neurodegeneration remains an unsolved question critical for the understanding of disease mechanism and the design of efficacious drugs.

**Methods:** Using a recombinant hyperphosphorylated tau protein (p-tau) synthesized by the PIMAX approach, we examined how cells responded to the cytotoxic tau and explored means to enhance cellular resistance to tau attack.

**Results:** Upon p-tau uptake, the intracellular calcium levels rose promptly. Gene expression analyses revealed that p-tau potently triggered endoplasmic reticulum (ER) stress, Unfolded Protein Response (UPR), ER stress-associated apoptosis, and pro-inflammation in cells. Proteomics studies showed that p-tau diminished heme oxygenase-1 (HO-1), an ER stress associated anti-inflammation and anti-oxidative stress regulator, while stimulated the accumulation of MIOS and other proteins. P-tau-induced ER stress-associated apoptosis and pro-inflammation are ameliorated by apomorphine, a brain-permeable prescription drug widely used to treat Parkinson’s disease symptoms, and by overexpression of HO-1.

**Conclusion:** Our results reveal probable cellular functions targeted by hyperphosphorylated tau. Some of these dysfunctions and stress responses have been linked to neurodegeneration in Alzheimer’s disease. The observations that the ill effects of p-tau can be mitigated by a small compound and by overexpressing HO-1 that is otherwise diminished in the treated cells inform new directions of Alzheimer’s disease drug discovery.

## Background

Alzheimer’s disease (AD) is the main cause of adult dementia afflicting more than 30 million patients world-wide. Except a small population of early-onset AD that are caused by mutations in APP, PSEN1 or PSEN2 leading to elevated synthesis of amyloid β (Aβ) peptides[1], most AD cases are sporadic and late-onset. In both familial and sporadic cases, a characteristic feature is the hyperphosphorylation of tau[2]. Tau is a microtubule associated protein[3] encoded by the MAPT gene. Despite the well-documented microtubule stabilizing activity in vitro, the physiological role of tau remains to be fully delineated[4]. For example, homozygous MAPT knockout mice developed normally without overt neurological phenotypes[5–7]. Older mice with tau deficiency do show motor and memory issues. However, these neurological defects appear to be dependent on the genetic background of the animals[8, 9]. In normal brain, tau phosphorylation is restricted to 2 - 3 phosphate per molecule[10]. Both the tau protein level and the degree of phosphorylation increase significantly in AD patients[11]. Because hyperphosphorylation disrupts the interaction between tau and microtubule[12], it has been suggested that the loss of function caused by abnormal phosphorylation contributes to the AD pathology. The relatively minor cognitive decline in tau knockout mice contradicts this model. Accumulating evidence instead suggests that a gain of toxicity of tau is a pivotal driver of the disease. For example, the spatiotemporal distribution of hyperphosphorylated tau in the form of paired helical filaments (PHFs), straight filaments (SFs), or neurofibrillary tangles (NFTs) robustly correlates with the progression of cognitive impairments[13, 14]. Cell culture and transgenic mouse models showed that reduction of tau markedly ameliorated the adverse effects resulting from Aβ accumulation[15, 16]. Finally, a cohort of neurodegenerative tauopathies share the trait of the deposition of hyperphosphorylated tau, wildtype or bearing a mutation, without Aβ plaques[17, 18]. Transgenic overexpression of the P301S mutant tau from frontotemporal dementia with Parkinsonism - chromosome 17 (FTDP-17)[19, 20] causes progressive accumulation of tau inclusions, accompanied by neuronal loss and brain atrophy in 9-12 months old mice[21–23]. Intrahippocampal injection of preformed fibrils of recombinant tau to young PS19 mice induced the formation and spread of inclusions closely resembling the AD neurofibrillary tangles[24]. Together, these reports demonstrated clearly that misfolded tau without Aβ is sufficient to drive neurological pathology in vivo.

Despite the increased interest in tau-mediated pathology that likely drives neurodegeneration in tauopathies, the mechanism by which the abnormally phosphorylated and misfolded tau inflicts cellular damage or death remains enigmatic. While neurofibrillary tangles are a conspicuous pathological character of AD and tauopathies, accumulating evidence argues against a causal role of these large tau conformers in neurodegeneration. Instead, the pre-tangle, soluble p-tau is likely the culprit. Human neurons bearing neurofibrillary tangles may live for decades[25]. In AD brains, neuronal loss in the superior temporal sulcus region exceeded the amounts of neurofibrillary tangles[26]. Tau knockout mice that overexpressed human tau developed neurofibrillary tangles, but the presence of such tau deposition did not correlate directly with cell death[27]. In rTg4510 mice where the expression of a human mutant tau is repressible, neuronal death was stopped upon repression of the human tau, yet neurofibrillary tangles continued to form[28], clearly separating tangle formation from neurotoxicity. Similarly, in a fly model that expressed pan-neuronally a mutant human tau, neurodegeneration was observed in the absence of discernible neurofibrillary tangles in the brain[29]. Instead of the well-structured neurofibrillary tangles, pre-tangle, soluble tau oligomers are increasingly recognized as a causal factor for neuronal loss in AD. Oligomeric tau from a synthetic source or from AD brains impaired membrane, and caused cell death[30–32]. The propagation of tau-mediated damages involves tau secretion[33, 34], uptake and internalization[35–38], and seeding conformational and functional changes of the endogenous tau[39–41]. The prion-like propagation of tau pathology appears to be the underlying mechanism for the progression of neurodegeneration[14, 42].

As to therapeutics development for AD and tauopathies, stopping the action of the pathological tau is a sensible yet comparatively under-developed approach. Many natural compounds reportedly reduced AD markers including Aβ and neurofibrillary tangles in animals[43–46]. Yet no success in clinical trials was reported. High-throughput screens for small molecule compounds that inhibited heparin-induced aggregation of unmodified tau have been reported as well[47–52]. However, a substantial portion of hits from these screens turned out to be non-specific redox modulators[53]. As such, the only tau aggregation inhibitor advanced to a phase 3 clinical trial failed to reach the primary end point[54–56]. One likely reason underlying these disappointing results is the use of unphosphorylated tau in an inducer-triggered aggregation assay. Disease tau is hyperphosphorylated. Heparin is a frequently used inducer for tau aggregation in vitro[57]. Heparin is an anticoagulant, and it remains to be verified as a pathophysiologically relevant factor for AD. Taking advantage of the inducer-free aggregation and cytotoxicity of hyperphosphorylated tau produced by the PIMAX approach[58, 59](see below), a pilot screen identified apomorphine and raloxifene[58], two prescription drugs shown to preserve the memory of transgenic mice[60] and to reduce the risks of dementia in humans[61], respectively. Besides small compounds, immunotherapy targeting p-tau has begun producing promising results. A recent report of the development of single-domain antibodies against pathological tau affords a novel direction for drug development[62].

PIMAX (<u>P</u>rotein Interaction <u>M</u>odules-<u>A</u>ssisted function <u>X</u>) is a versatile recombinant DNA technology that enables the synthesis of proteins in E. coli with a desired post-translational modification[59]. The full-length hyperphosphorylated tau (referred to as p-tau hence forth) produced by PIMAX bears phosphorylation marks that recapitulate many of those found associated with AD including phosphoepitopes used for AD staging on human postmortem samples, such as p-T181, p-S202, and p-T217[30, 63, 64], therefore linking this recombinant tau to the disease. Furthermore, without detectable tangle-like structures, PIMAX p-tau is prone to further aggregate into cytotoxic species in the absence of an artificial inducer such as heparin[30]. This cytotoxicity results from the activation of apoptosis and of the accumulation of mitochondrial superoxide, and was efficaciously prevented by several blood-brain-barrier permeant compounds such as apomorphine and raloxifene[58], hence paving a way to tau toxicity-based drug discovery. Conversely, compounds such as selective benzodiazepines used widely for a variety of neurological issues were found to enhance the aggregation and cytotoxicity of PIMAX p-tau[58], suggesting that certain risk factors for AD and AD-related dementia can be identified via the use of PIMAX p-tau[65, 66].

In addition to the action of a pathological tau, cellular functions crippled by tau also seem to be viable candidates for measures that aim to improve neuronal functions. Delineating how brain cells die in AD therefore may contribute significantly to the development of efficacious drugs. Of the many candidates, of special interest is the endoplasmic reticulum (ER) stress and Unfolded Protein Response (UPR). In eukaryotic cells, the ER serves many specialized functions including folding and assembly of membrane and secretory proteins, calcium storage and flux, and production of lipids and sterols[67]. As a compartment responsible for protein folding and many posttranslational modifications, the ER is exquisitely sensitive to alterations in proteostasis. High demands for protein folding and clearance of aberrant proteins or abnormal increase in protein synthesis disrupts ER homeostasis, and causes ER stress[67] that triggers the UPR to clear those unfolded and misfolded protein. Prolonged, unresolved ER stress leads to apoptosis and cell death. Given the conspicuous presence of misfolded Aβ and tau in AD, it is not surprising that postmortem samples of AD showed elevated ER stress and UPR[68–70]. The UPR classically encompasses the activation of three separate ER transmembrane stress sensors, inositol-requiring enzyme 1α (IRE1α), protein kinase (PKR)-like endoplasmic reticulum kinase (PERK) and activating transcription factor 6 (ATF6) [67, 71]. Among them, PERK is activated to phosphorylate the eukaryotic translation initiation factor eIF2α and thus attenuates global protein synthesis to reduce the luminal client load in response to ER stress. However, under prolonged ER stress or acute injuries to the ER, PERK-mediated UPR induces proapoptotic transcription factors, including ATF3, ATF4, and CHOP (C/EBP homologous protein), as well as BH3-only BCL2 family members, such as NOXA and BIM[67, 72, 73] CHOP can induce cell cycle arrest or apoptosis by regulating expression of multiple genes encoding proapoptotic factors including DR5 (death receptor 5), TRB3 (Tribbles homolog 3), CAVI (carbonic anhydrase VI)[73, 74]. CHOP also contributes to apoptosis through activating ERO1α, an ER oxidase that promotes hyperoxidization of the ER[75]. Additionally, ER stress, UPR, and inflammatory responses are interconnected through various mechanisms, including the production of reactive oxygen species (ROS), the release of calcium from the ER, the activation of the transcription factor nuclear factor-κB (NF-κB) and a mitogen-activated protein kinase JNK (JUN N-terminal kinase)[72]. ER stress response promotes production of proinflammatory cytokines in macrophages, fibroblasts, astrocytes, or epithelial cells associated with autoimmune, metabolic, and neurodegenerative diseases[67].

The chronic presence and accumulation of Aβ and p-tau in the brain underlie the onset and progression of AD neurodegeneration. Such misfolded proteins are very likely to contribute to the etiology through the activation of ER stress and unfolded protein response. Here we show that p-tau causes cell death likely through the activation of ER stress, UPR, and inflammatory response. In addition, molecular changes known to activate these stress responses, such as dyshomeostasis of intracellular calcium and damages in cytoskeleton, are among those early responses following the treatment of p-tau. These experimental results shed light on a critical molecular mechanism of tauopathy.

## Methods

### Materials

Chemicals were purchased from Sigma unless indicated otherwise. Synthetic oligonucleotides were purchased from Integrated DNA Technologies, Inc. (Coralville, IA). Lipofectamine™ 2000 and Fluo-4 AM (F14201) were purchased from Invitrogen. Opti-MEM, Gibco DMEM/F12 (10565018) was purchased from Thermo Fisher Scientific. 150-mesh copper square grids (100489-712) were from VWR. The DA9 tau antibody was a generous gift of Dr. Peter Davies [76]. Plasmid pCAG-EGFP was a generous gift of Dr. Ralston. Plasmid pCX-HO1-2A-EGFP was ordered from Addgene. Goat-anti-Mouse IgG+IgM (10 nm gold) was a generous gift of Dr. Alicia Withrow. Cell Counting Kit-8 (CCK-8) (GK10001) was purchased from GlpBio. Alexa Fluor 594 protein labeling kit (A10239) was purchase from Invitrogen. The filamentous actin (F-actin) probe, Alexa Fluor 488 phalloidin (A12379), was purchased from ThemoFisher Scientific.

### Primary mouse cortical neuron culture and in vitro cytotoxic assay

Mouse cortical neurons were prepared from E16.5 mouse embryos. The cortex was mechanically dissociated by passing through a glass pipette for twenty times and then filtered through a 70-μm nylon mesh filter (BD Biosciences). Cells were plated onto plastic culture plates coated with 30 μg/ml poly-L-lysine and 2 μg/ml laminin and cultured in Neurobasal Medium containing 10% FBS, 100 U/ml penicillin and streptomycin, and N2 supplement and B27 supplement (Invitrogen). The cells were kept at 37 °C in a humidified atmosphere of 5% CO2 and 95% air. p-tau and Apomorphine (Cayman, Cat. #16094) were added into the culture medium at 6 D.I.V. cortical neurons. The Cell viability of 10 D.I.V. or 13 D.I.V. cortical neurons was assayed by a colorimetric procedure with cell counting kit-8 (Dojindo Laboratories, E-CK-A362) according to the manufacturer’s instructions.

### Transmission electron microscopy and immunogold staining

10 µM p-tau was incubated with or without compounds in 20 mM Tris buffer (pH7.4) at 37 °C for indicated time. The sample was 10-fold diluted and incubated with 2.5% glutaraldehyde for 5 min. Then 20 µL of the sample was fixed on a 150-mesh copper square grid and negatively stained by 1% uranyl acetate for 10 seconds. The sample was imaged by JOEL 1400 Flash TEM. For immunogold staining, 20 µL of the sample was fixed on the grid with 2.5% paraformaldehyde for 5 minutes and blocked with 1% BSA at room temperature for 1 hour. Then, the grid was incubated with primary antibody (DA9 antibody 1:1000) at 4 degrees overnight followed by secondary 10 nm gold antibody (1:50) at room temperature for 2 hours. The grid was then incubated with 2.5% glutaraldehyde for 15 min and stained by 1% uranyl acetate for 10 seconds. Then the sample was imaged by JOEL 1400 Flash TEM.

### Transfection

pCAG-EGFP or pCX-HO1-2A-EGFP (Addgene, #74672) were transfected using a standard Lipofectamine 2000 protocol. SH-SY5Y cells were seeded with 2 x 105 cells/ml and incubated for overnight for transfection. Plasmid/Lipofectamine 2000 mixture ratio of 0.8 μg DNA: 1.0 μL lipofectamine 2000 in Opti-MEM were added into the cells which were changed to serum-free media. Cultured for 6 hours and the media were refreshed to the media with 5% FBS and refreshed again to the media with 10% FBS after 18 hours. The cells were passaged for amplification at 24-hour post-transfection if needed. To rule out the interference from non transfected cells and get viable transfected cells for corresponding experiments, DAPI-negative and GFP-positive cells were sorted with BD influx cell sorter.

### Western Blotting

To make extract for western procedure, cells were rinsed with PBS and scraped off the plates. The cell pellet was collected after spined down with 2000 rpm for 10 mins. The pellet was lysed using Sherr Lysis Buffer containing 50 mM HEPES pH 7.5, 150 mM NaCl, 1 mM EDTA, 2.5 mM EGTA, 0.1% Tween-20, 10% glycerol, 50µM PMSF, 1mM Benzamidine, 1mM Sodium Bisulfite, 1mM DTT. The supernatant was quantitated by a Bradford Protein Assay Kit (Thermo Fisher Scientific). 150 µg protein lysates were separated by SDS-PAGE and transferred on to polyvinylidene difluoride (PVDF) membranes. Membranes were blocked with 5% milk dissolved in 1X PBS; pH 7.4 with 0.1% Tween-20 for 1h in room temperature. Then, membranes were incubated with relevant antibodies at 4°C overnight followed by HRP–conjugated secondary antibody. The proteins were visualized by ECL reagent (Thermo Fisher Scientific). Primary antibodies: MIOS (# 13557S, Cell Signaling), HO-1 (ab13248, Abcam) were diluted in 5% milk ratio of 1/1000, and 1/1000, respectively. Secondary antibodies were diluted in 5% milk ratio of 1/5000.

### CCK-8 cell viability assay

To measure the cytotoxicity of p-tau, Cell Counting Kit-8 (CCK-8) assay was used. SH-SY5Y cells were seeded in 96-well plate with the density of 2 x 10 cells/ml in 100 µL DMEM/F12 media and grown for overnight. Then, cells were incubated with p-tau for overnight. The cell viability was assessed by adding 10 µL of CCK8 solution to the cells at specific time points. Hereafter, the cells were incubated for 2 hours at 37°C before measurement of the OD values at 450 nm using a BioTek microplate reader. Data were normalized by the mean value of the no-treatment group.

### Quantitative real-time PCR

Total RNAs from SH-SY5Y neuroblastoma cells were extracted using Trizol by following manufacture’s instruction and complementary DNA was synthesized from 500 ng of total RNA to with High Capacity cDNA Reverse Transcription Kit (AB applied biosystems). The abundance of mRNA was measured by quantitative real-time PCR analysis using the SYBR Green PCR Master mix (Applied Biosystems). PCR was carried out with 96-well plate using Applied Biosystems 7500 Real-Time PCR Systems. Mouse β-actin was used reference gene. The sequences of the primer sets were listed in S-Table 1.

### Fluorescence microscopy, ER and cytoskeleton staining

SH-SY5Y cells were seeded to 24-well plates or 2-chamber coverglass (Lab-Tek) at the density of 4 x 10 cells/ml in media (DMEM/F12, 10% FBS, pen/strep) and cultured for 24 hours at 37 °C, 5% CO_2_. These cells were then treated with varying amounts of p-tau or Alexa conjugated p-tau with or without compounds for the indicated time. For calcium imaging, 1 µM of Fluo-4 AM was added to the cells 1 hour prior to the imaging for free calcium staining. The live cells were imaged by Nikon A1 confocal microscope, Olympus FluoView Filter confocal microscope or Biotek Cytation 5 Cell Imager.

For actin staining and quantification, cells were incubated with Alexa Fluor 488 phalloidin (Thermo Fisher), a high-affinity filamentous actin (F-actin) probe conjugated to green-fluorescent Alexa Fluor 488 dye for 1 h. The stained cells were mounted with DAPI containing the ProLong Glass Antifade Mountant (ThermoFisher), and observed using Cytation Confocal Imaging Reader (Biotek/Agilent). For ER staining, SH-SY5Y cells cultured in slide chambers were incubated with 1 μM of ER-Tracker Red (Thermo Fisher) for 20 min at 37°C according to manufacturer’s instruction. The stained cells were mounted with Prolong Glass Antifade Mountant, and observed using Zeiss fluorescent microscopy.

For ER staining, SH-SY5Y cells cultured in slide chambers were incubated with 1 μM of ER-Tracker Red (Thermo Fisher) for 20 min at 37°C according to manufacturer’s instruction. The stained cells were mounted with Prolong Glass Antifade Mountant, and observed using Zeiss fluorescent microscopy.

ImageJ software was utilized to quantitatively analyze fluorescent intensities. Cells of interest and regions of interest (ROI) were selected and measured for mean fluorescence intensity. A region next to the cell or region of interest that has no fluorescence was selected for background fluorescence measurement. The above-mentioned steps for the other cells in the field of view were repeated. Corrected Total Cell Fluorescence (CTCF) were calculated based on: CTCF = Integrated Density – (Area of Selected Cell x Mean Fluorescence of Background readings)

### LC-MS/MS

Three micrograms of each trypsin digested proteins were analyzed by nano LC-MS/MS with a Waters M-Class HPLC system interfaced to a ThermoFisher fusion Lumos mass spectrometer. The MS was operated in data-dependent mode, with the Orbitrap operating at 60,000 FWHM (full width at half maximum) and 15,000 FWHM for MS and MS/MS respectively. Advanced Peak Determination (APD) was enabled, and the instrument was run with a 3s cycle for MS and MS/MS for 4 hrs. Data were processed with MaxQuant version 1.6.14.0[77].

### Data analysis and statistics

Data are presented as mean ± SEM (standard errors of the mean). Statistical comparisons and analyses between 2 groups were performed by two-tailed unpaired Student’s t test. For 3 or more groups, one-way ANOVA analysis test were used for parametric comparisons. A p < 0.05 was considered as statistical significance.

## Results

### P-tau produced by PIMAX confers toxicity to primary neurons

We previously reported that the p-tau (p-tau) produced by PIMAX exhibited significant cytotoxicity to HEK293T and SH-SY5Y cell lines, and to mouse neurons differentiated from neural stem cells of prenatal mice[30]. In addition to significant overlapping with disease related phosphorylation sites such as T181 and T217[63, 78], the stoichiometry of phosphorylation (estimated to be 11 – 14 phosphate per tau molecule, Supplemental Figure 2) also is consistent with that in the disease[10]. To further ascertain the neurological relevance of the recombinant p-tau, we isolated primary neurons from mouse fetuses (E16, Figure 1A) and tested their survival upon treatment of increasing concentrations of p-tau. As anticipated, clear p-tau dose-dependent viability loss of primary neurons was observed (Figure 1B). Intriguingly, the post-mitotic primary neurons are significantly more resistant to p-tau, with the estimated LD_50_ (concentration of p-tau causing 50% cell death) to be between 2 and 4 µM. In comparison, the LD_50_ for SH-SY5Y cells, HEK293T cells, and mouse neurons differentiated from neural stem cells are 0.57 µM, 0.75 µM, and 1.1 µM, respectively[30]. The pre-tangle soluble tau in the frontal cortex of AD patients has been estimated to be between 4 and 7 µM[10]. The LD_50_ of p tau against the primary mouse neurons agrees well with the pathophysiological concentration of disease tau.

**Figure 1.**
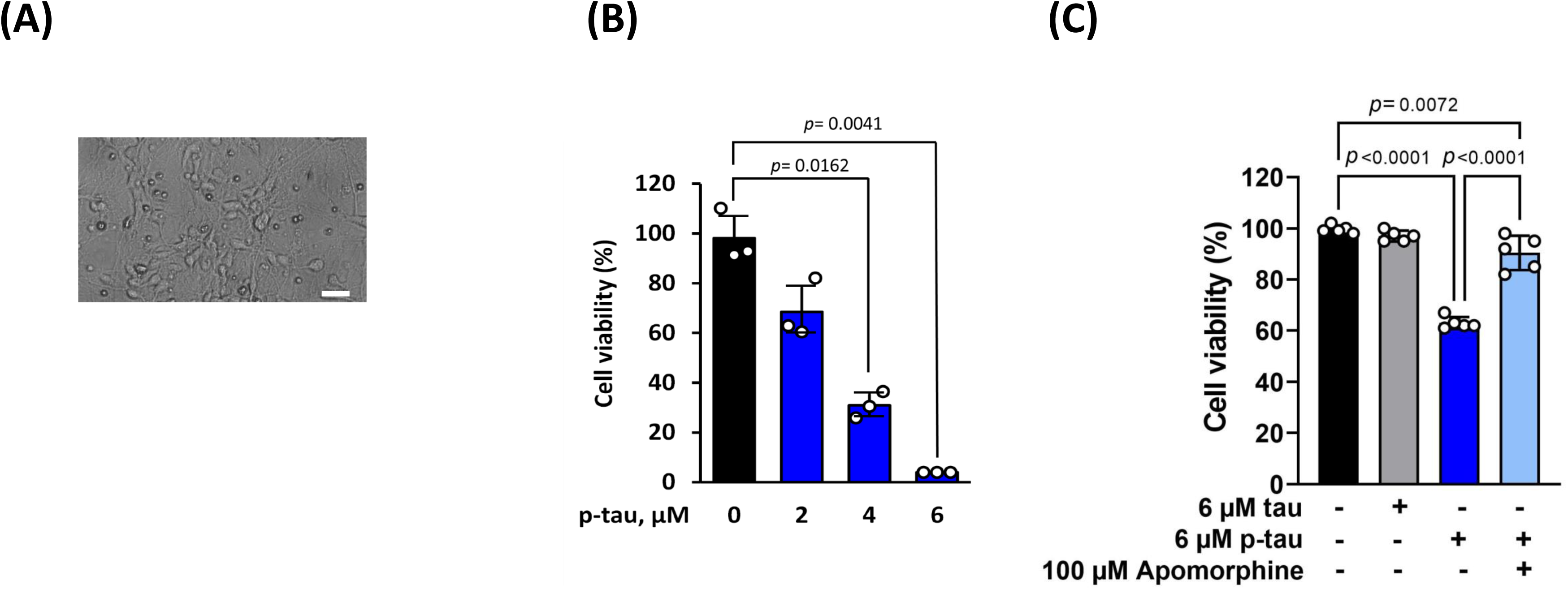
P-tau confers cytotoxicity in mouse primary neurons. (A) A representative image of primary E16.5 mouse cortical neurons cultured at 6 D.I.V. Scale bar represents 13.8 μm. (B) Primary cortical neurons at D.I.V 6 were treated with p-tau for 7 days. CCK-8 assay was performed to measure the cell viability. N = 3 wells per group. (C) Cortical neurons from the E16.5 mouse cortex were cultured and treated with tau or p-tau. Apomorphine was included in one of the two p-tau treatment samples. Treatments started at 6 D.I.V. Neurons were harvested at 10 D.I.V. for viability assay by cell counting kit-8. Data from five independent experiments were shown as means ± SD, one-way ANOVA with Tukey’s post hoc test.

We further explored a character of p-tau that might be of critical significance in AD disease mechanism and drug discovery, that is, amelioration of the phosphorylation-dependent neurotoxicity of tau by apomorphine, a prescription drug previously found in our pilot screen to inhibit p-tau cytotoxicity in vitro[58]. Compared with p-tau that effectively reduced the viability of primary neurons, the unphosphorylated tau (tau) imposed no discernible effect (Figure 1C). Intriguingly, the neurotoxicity of p-tau was quantitatively prevented by apomorphine[58].

Data shown in Figure 1 are in excellent agreement with the observations made with non-neuronal cells in our previous reports[30, 58], i.e., the hyperphosphorylation-dependent cytotoxicity of tau is amendable to apomorphine inhibition. We suggest that p-tau may target a pathway(s) that is shared by many different cell types, and that pharmacological control of this pathway(s) may lead to the discovery of novel, efficacious AD drugs targeting the pathogenicity of abnormally phosphorylated tau.

### Modulation of pre-PHF conformational changes of p-tau by apomorphine and raloxifene

To gain further insights into p-tau toxicity and its control, we examined the time-dependent alterations of p-tau structures by electron microscopy. That non-neuronal cells lost viability within one day of p-tau treatment argued against the involvement of p-tau with significantly ordered structures, for amyloid fibrils typically required days of growth under such conditions as agitation and in the presence of an inducer or seeds. Indeed, results from animal models argue against a direct pathogenic role of neurofibrillary tangles[29, 79]. Instead, tangles may protect neurons by sequestering the otherwise diffusible and toxic tau species[80]. Consistently, the cytotoxicity of p-tau produced by PIMAX increased in the first three days during an inducer-free aggregation reaction, but decreased progressively afterwards[30], suggesting that the cytotoxic species might be an intermediate before evolving into a less toxic PHF and NFT state. We therefore hypothesized that the cytotoxic p-tau existed as structures distinct from the terminal neurofibrillary tangles or paired helical filaments. Furthermore, because the cytotoxicity of p-tau is effectively diminished by apomorphine, we further hypothesized that the structures of p-tau would be amenable to modulations by apomorphine. To test these hypotheses, we performed transmission electron microscopy (TEM) to examine whether and how p-tau aggregates might change their overall sizes and morphology over five days of incubation. TEM results shown in Figure 2 and immunogold staining shown in Supplemental Figure 3 concurred with our notions.

**Figure 2.**
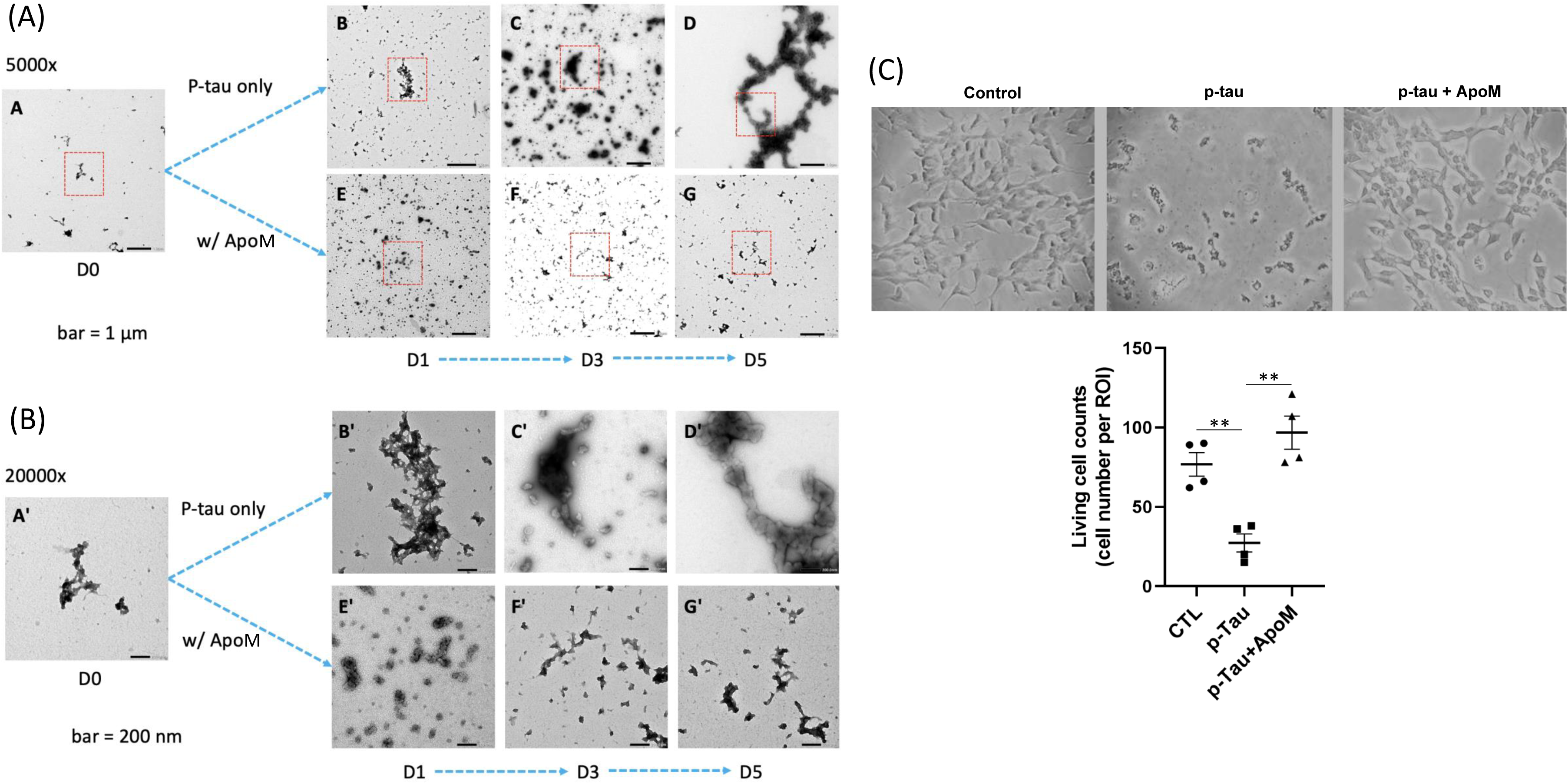
Cytotoxic p-tau (p-tau) forms amorphous aggregates but not tangle-like or PHF structure in vitro, while apomorphine affects the progression of p-tau aggregation and cytotoxicity. (A) P-tau sampled from 0, 1, 3, and 5 days of in vitro aggregation reactions with or without apomorphine. 1 µM of the sample was negatively stained and observed by TEM. A – D: p-tau only; E – G: p-tau aggregation in the presence of apomorphine (E – G). Scale bar = 1 µm. (B) Higher magnification of boxed areas in A shows detailed structural changes of p-tau with or without the influences of apomorphine. Scale bar = 200 nm. (C) Apomorphine protected cells from the cytotoxicity of p-tau . (C-a) Cell morphology of SH-SY5Y cells under the treatment of vehicle, p-tau (0.5 µM), or p-tau (p-tau) with apomorphine (ApoM, 5µM) for 20 hr, was visualized by bright field and phase contrast microscopy. Magnification: 20X. (C-b) SH-SY5Y cell counts after the treatment of vehicle, p-tau, or p-tau plus ApoM. Mean ± SEM (n = 4 biological repeats). ** p ≤ 0.01.

Without an aggregation inhibitor, p-tau assembled gradually into larger ensembles with minimal resemblance to paired helical filaments after five days of incubation (panels A – D). Panel A (5,000x magnification) images show the overall distribution of p-tau structures that were mostly amorphous and fractal, consistent with our prior report[30]. Panel B (20,000x magnification) images reveal details of boxed regions in panel A. There were occasional small clusters (<30 nm in diameter) and larger fractals (panels B and B’). These clusters grew bigger and more ordered. Many ∼50 - 100-nm complexes were seen in day-3 samples (see panels C and C’). From day 3 to day 5, these structures further conglomerated to form much larger and longer aggregates (panels D and D’). However, these did not resemble the canonical fibrillar structures seen in PHF and NFT. Multiple TEM observations of separate batches of p-tau consistently showed this trend with few, if any, PHF-like fibrils. These results were consistent with the fact that tau is an intrinsically disordered protein that may progress through different structural propensities[81] before forming the final species of PHFs and NFTs. It is possible that a portion of p-tau molecules were irreversibly trapped as amorphous deposits in vitro and in vivo[82].

In the presence of a p-tau aggregation inhibitor, apomorphine, the evolution of p-tau conformers deviated significantly. Apomorphine prevented the formation of the larger, more structured entities (compare panel D with panel G, Figure 2) seen in day 5 of the p-tau-alone sample. Apomorphine first organized p-tau into numerous globular structures that were about 30 - 50 nm in diameter in the first 24 hours of aggregation. These globular structures organized into larger arrays of ∼200 nm in length, but eventually were dismantled into small particles with no clear features in day 3 and day 5 (panels F, and F’). In addition to apomorphine, a prescription drug called raloxifene (under the market name of Evista) was also found in our previous drug screen to be an effective inhibitor for the aggregation and cytotoxicity of p tau[58]. Parallel TEM assays were conducted with raloxifene. The effects of raloxifene on the structures of p-tau aggregates can be seen in the micrographs (Supplemental Figure 4). Though apomorphine and raloxifene directed p-tau toward somewhat different paths of folding, they both impacted significantly on the growth of p-tau complexes, consistent with their ability to inhibit p-tau aggregation[58] and cytotoxicity.

The above TEM results revealed the early structural development of p-tau that can be physically modulated by apomorphine and by raloxifene. When treated with 0.5 µM of p-tau for 20 hours, the neuroblastoma SH-SY5Y cells showed significant cell shrinkage and death (Figure 2C and Supplemental Figure 4). Such drastic morphological abnormalities were prevented to a great extent by both apomorphine (Figure 2C) and raloxifene (Supplemental Figure 4, bottom). However, some cells formed small clusters. Fluoresceine diacetate (FDA) and propidium iodide (PI) differential staining confirmed that most of the p-tau-treated cells were kept alive by apomorphine or raloxifene, but down to approximately 40% viable without these compounds (data not shown, and see reference [30]).

In summary, Figures 1 and 2 show that p-tau is able to inflict significant damages to SH-SY5Y and primary neurons in a fashion that is quantitatively prevented by apomorphine. While not yet tested, it is likely that raloxifene would exhibit a comparable cytoprotective function for primary neurons as well. To deconvolute the cytotoxicity of p-tau, the rest of this work aimed to delineate the sub-cellular responses following p-tau treatment.

### Hyperphosphorylated tau targets multiple cellular functions

Understanding how p-tau impacts cell function and viability will help promote the drug discovery endeavor. Since p-tau exhibits cytotoxicity of all cells that we have tested so far (HEK293T, SH-SY5Y, and neurons from neural stem cells or mouse fetuses) in a manner that is averted by apomorphine, we suspect that the fundamental mechanism by which p-tau kills cells is shared by these cells. Due to the relatively straightforward culturing procedures, we therefore used the neuroblastoma cell line SH-SY5Y as the surrogate for the subsequent mechanistic studies. To this end, we first examined whether cells could absorb recombinant p tau. A sub-lethal dose (0.1 µM) of p-tau labelled with the Alexa594 fluorescent tracer was added to the SH-SY5Y neuroblastoma cells and incubated for 16 hours. Confocal microscopy showed that majority of cells remained healthy, and that p-tau indeed entered the cells, as indicated by the diffused cytoplasmic Alexa594 signals (Figure 3A). Furthermore, tau formed many puncta, consistent with the tendency of p-tau molecules to aggregate into oligomers as evidenced in Figure 2. An alternative explanation is that certain surface receptors may be bound by p-tau[33, 35]. We were also curious to know whether apomorphine protected cells from p-tau by preventing the uptake of this protein. To our surprise, despite the clear power of modulating the higher-ordered structure of p-tau, apomorphine did not seem to cause appreciable diminishment of p-tau absorption (Figure 3B). This result strongly suggested that the major cellular damages caused by p-tau were inside the cells, not the membrane integrity.

**Figure 3.**
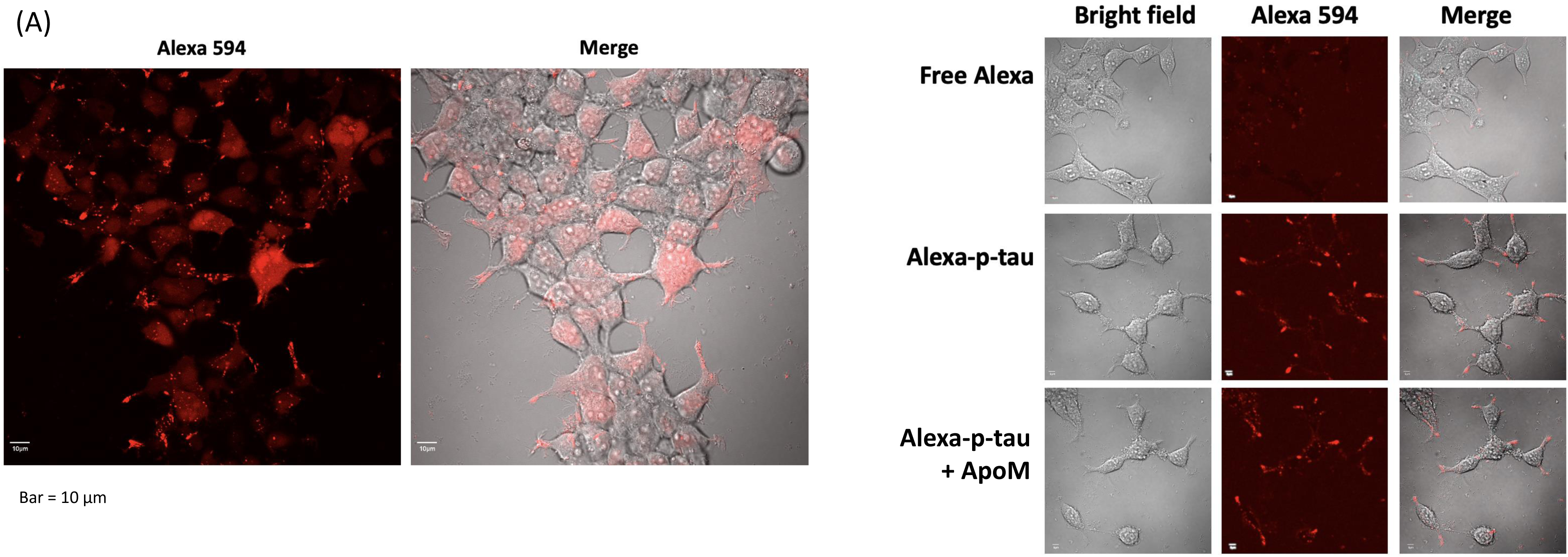
Confocal microscopy reveals the p-tau (p-tau) uptake by cells but not blocked by apomorphine. (A) SH-SY5Y cells were treated with 0.1 µM Alexa 594-conjugated p-tau for 16 hours. Cells were imaged by the Olympus FV microscope, 20X objective. (B) Apomorphine does not inhibit appreciably the uptake of p-tau by SH-SY5Y cells. Cells were treated with 0.1 µM Alexa 594-conjugated p-tau in the presence or absence of 1 µM apomorphine for 20 hours before imaging the Olympus FV microscope, 40X objective. Note the p-tau-dependent absorption (middle row) was not affected by apomorphine (bottom row).

AD is a proteinopathy, i.e., a protein misfolding disease[83]. The first line of defense against misfolded proteins is the endoplasmic reticulum. ER plays major roles in responding to dyshomeostasis by activating ER stress response or UPR[67] and, if the burden exceeds the repair capacity, triggers ER stress-associated pro-apoptotic actions. To understand whether p tau inflicted cytotoxicity via ER stress and UPR signaling pathways, SH-SY5Y cells were treated with 0.5 µM of p-tau for 24 hours. Quantitative real-time PCR (qPCR) was then performed with the total RNAs isolated from the treated cells to profile the expression of the genes encoding ER stress markers and major components of UPR signaling, ER stress-associated apoptosis, as well as pro-inflammatory responses. In response to p-tau challenge, expression levels of the transcripts encoding the ER chaperone BiP/GRP78, the primary UPR transducer IRE1α, the UPR transcriptional activator spliced XBP1 (XBP1s), as well as the UPR targets, including P58ipk and ERO1, were significantly increased (Figure 4A). The expression levels of the transcripts encoding major mediators or markers in ER stress-associated pro-apoptotic pathways, including ATF4, ATF3, DR5, TRB3 and GADD34, also elevated drastically (Figure 4B). Moreover, p-tau stimulated inflammatory pathways in SH-SY5Y cells, as evidenced by the heightened expression of the genes encoding the inflammatory cytokines TNFα, IL8, and INFγ in response to the challenge from p-tau (Figure 4C). Importantly, in the presence of apomorphine, the levels of all mediators or markers of ER stress, ER stress-associated apoptosis, and inflammatory responses induced by p-tau were at or near the no-treatment controls (Figure 4A - C, gray bars). The dramatic suppressive effects of apomorphine on tau-induced ER stress-associated apoptotic and inflammatory responses provided an important clue for the cellular pathways pivotal to AD pathogenesis.

**Figure 4.**
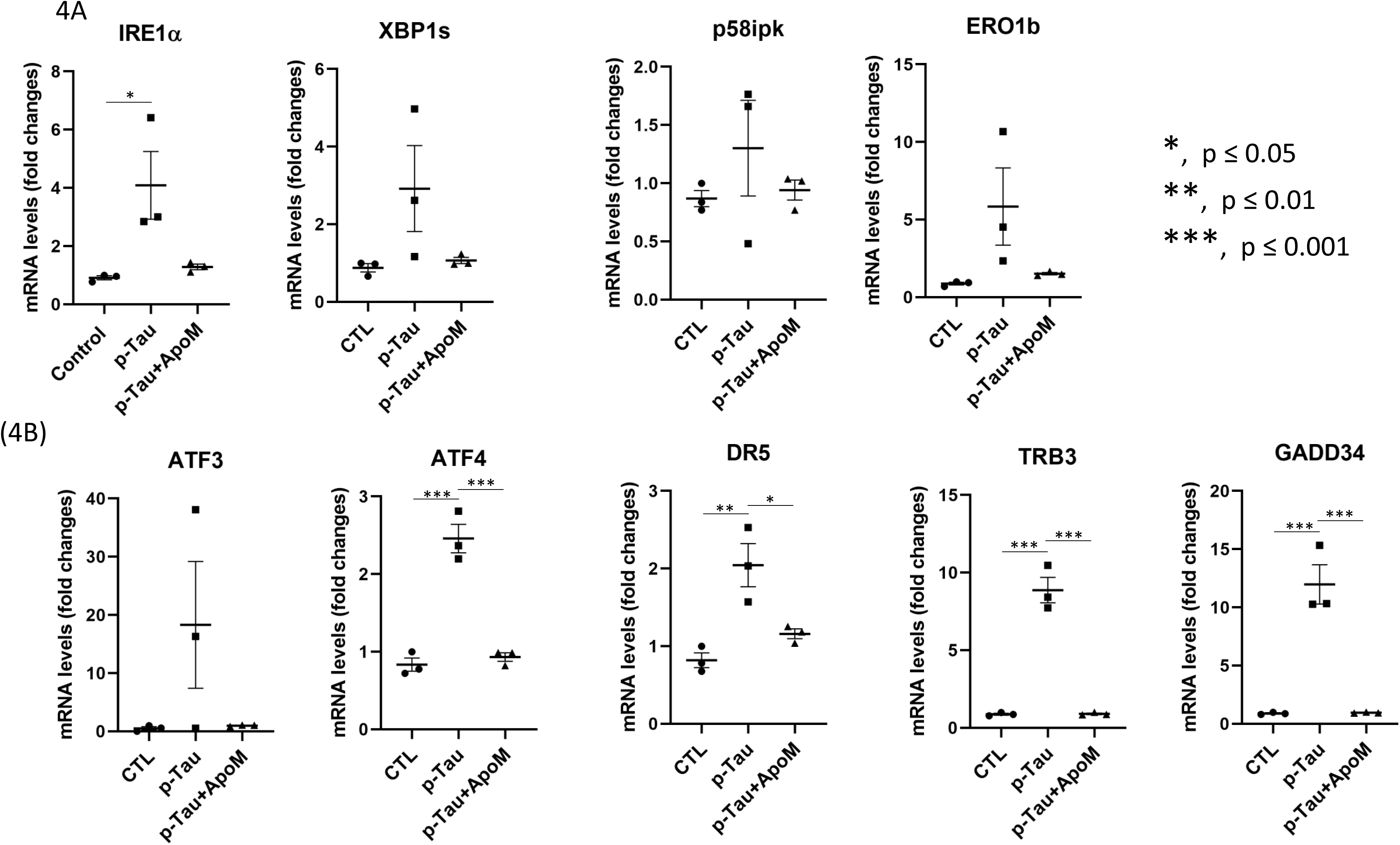
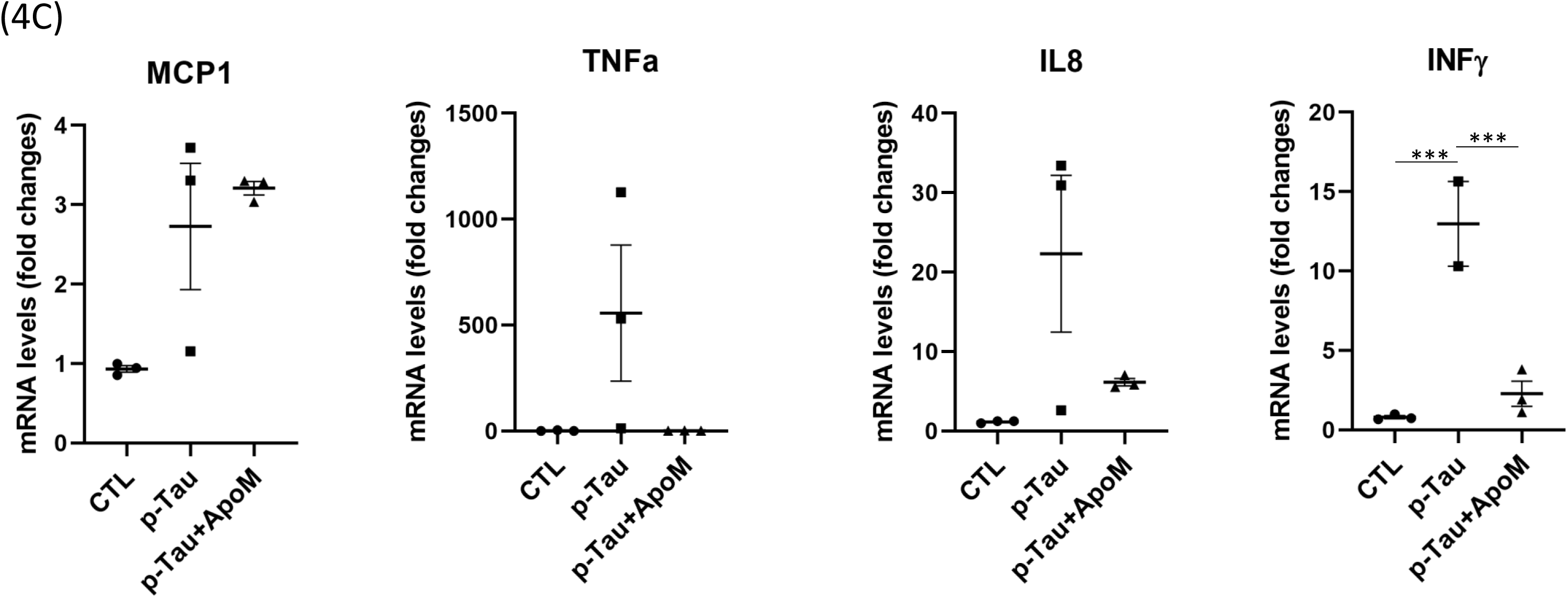
P-tau activates Unfolded Protein Response, ER stress-associated apoptosis, and pro inflammatory responses that are ameliorated by apomorphine. (A) SH-SY5Y cells were treated with vehicle, p-tau (0.5 µM) without or with apomorphine (5 µM) for 20h. Quantitative real time PCR (qPCR) analysis of the transcripts encoding major ER stress markers or UPR mediators, including IRE1α, spliced XBP1 (SBP1s), P58ipk, and ERO1β in SH-SY5Y cells after the treatments. Mean ± SEM (n = 3 biological repeats). * p ≤ 0.05. (B) qPCR analysis of the transcripts encoding major mediators of ER stress-associated apoptosis, including ATF3, ATF4, DR5, TRB3, and GADD34 in SH-SY5Y cells treated with vehicle, p-tau, or p-tau plus apomorphine (ApoM) as described in panel A. Mean ± SEM (n = 3 biological repeats). * p ≤ 0.05; ** p ≤ 0.01; *** p ≤ 0.001. (C) qPCR analysis of the transcripts encoding representative mediators of pro-inflammatory response, including MCP1, TNFα, IL8, and INFγ in SH-SY5Y cells treated with vehicle, p-tau, or p-tau plus ApoM as described in panel A. Mean ± SEM (n = 3 biological repeats). *** p ≤ 0.001.

To rule out that apomorphine acted as a general suppressor of ER stress, we treated cells with tunicamycin, a classical ER stress-inducing reagent that can cause accumulation of unfolded or misfolded proteins in the ER by disrupting protein glycosylation[84]. In stark contrast to p-tau, tunicamycin-induced ER stress, ER stress-associated apoptosis, and inflammatory responses all remained high despite the co-treatment of apomorphine (Figure 5). Intriguingly, for an unknown reason, the effect of tunicamycin was enhanced by apomorphine without tau. Together, these results demonstrated that apomorphine is a specific inhibitor for the toxicity of p-tau.

**Figure 5.**
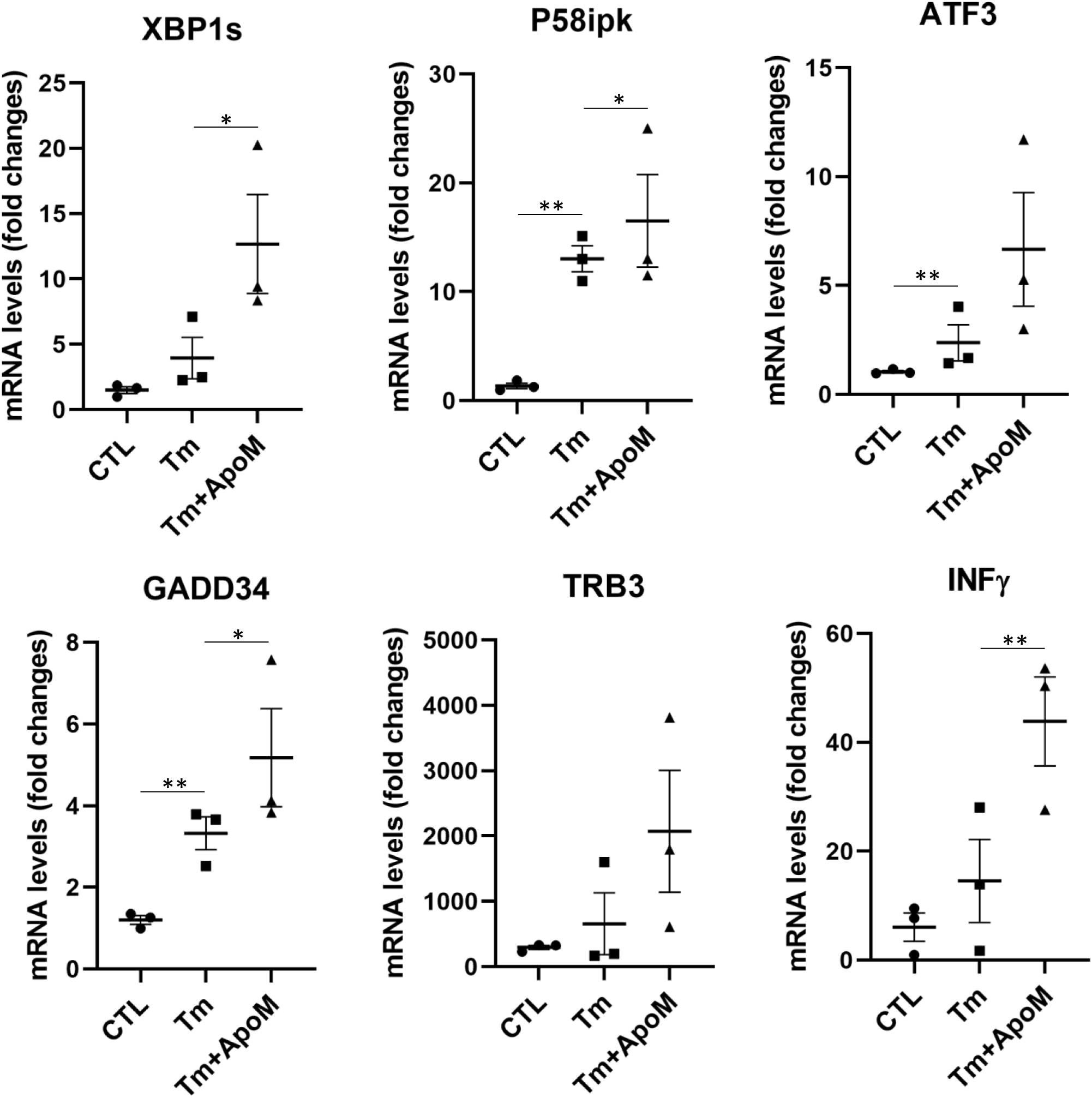
Apomorphine specifically antagonizes p-tau-, but not tunicamycin-induced ER stress. qPCR analysis of the transcripts encoding representative mediators of UPR, ER stress-associated apoptosis, and pro-inflammatory response, including XBP1s, P58ipk, ATF3, GADD34, TRB3, and INFγ in SH-SY5Y cells treated with vehicle, tunicamycin (Tm, 5µg/ml), or Tm (5µg/ml) plus ApoM (5µM) for 20h. Mean ± SEM (n = 3 biological repeats). * p ≤ 0.05; ** p ≤ 0.01.

### P-tau causes systemic changes in cells

How was ER stress activated by p-tau? Were there additional tau-inflicted impairments in these cells? The low concentration of p-tau used in assays shown in Figure 4 (≤ 1 µM, ∼0.005% wt/vol) did not appear to be sufficient to elicit UPR directly, for in the normal cortex, tau is estimated to be about 2 µM; and in the AD brain, this level increased to 8 – 10 µM[11]. We suspected that p tau triggered a systemic crisis that led to the activation of ER stress response. To test this hypothesis, we first examined the integrity of the cytoskeleton in p-tau-treated cells. The ER and the microtubule cytoskeleton form a co-extensive, dynamic system in the cytoplasm of the cells, and disruption of cytoskeleton can damage the structure of the ER and impair its function[85, 86]. Actin staining with phalloidin revealed a substantive destruction of intracellular cytoskeleton in SH-SY5Y cells when treated with p-tau (0.5 µM) for 24 h (Figure 6, top row). In contrast, SH-SY5Y cells displayed intact cytoskeleton structure when co-treated with p-tau and apomorphine, indicating that apomorphine was highly effective in protecting cytoskeleton from the insults of p-tau. Despite the massive cytoskeleton reorganization upon p tau challenge, the signals of ER were only modestly changed (middle row), implicating a functional ER that mediates robust UPR signaling in response to p-tau. Further, the presence of apomorphine did not alter ER abundance in the p-tau-treated SH-SY5Y cells. Additionally, there was no discernible change in the mitochondrial morphology in SH-SY5Y cells after the overnight treatment (bottom row).

**Figure 6.**
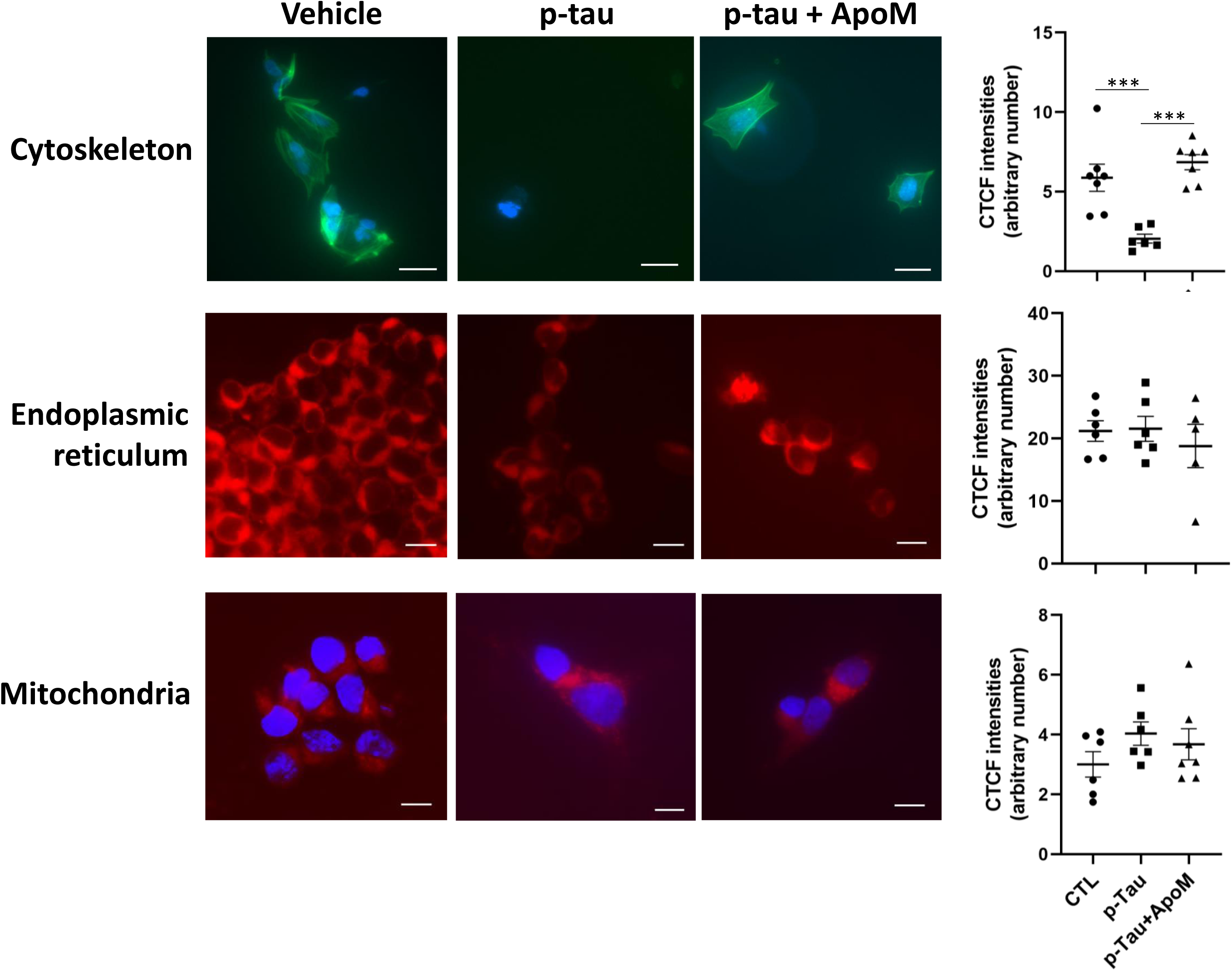
Apomorphine prevents p-tau-induced damages to cytoskeleton and ER, whereas mitochondria remained morphologically normal in all conditions. SH-SY5Y cells underwent the indicated treatment for 20 hours were examined for the integrity of cytoskeleton (top row), ER (middle row), and mitochondria (bottom row). Cell nuclei were counter-stained with DAPI (blue). Magnification: 60X. Scale bar: 20 µm. Intracellular cytoskeleton, ER, and mitochondria abundance, reflected by Corrected Total Cell Fluorescence (CTCF), were quantified. Mean ± SEM (n = 6 biological repeats). *** p ≤ 0.001.

We then examined the intracellular calcium flux as an early response to p-tau treatment. Calcium dyshomeostasis is associated with neurodegeneration[87]. A recent report demonstrated the connection between tau pathology and calcium dysregulation in aged macaques[88]. P-tau-treated cells showed mitochondrial superoxide accumulation and apoptotic activity, with the latter being suppressed by pre-treating cells with ROS scavengers such as Trolox and TEMPOL[30]. Both stresses might be explained by Ca^2+^ imbalance[89, 90]. To test this hypothesis, we used a cell permeant Ca^2+^ indicator, Fluo-4 AM[91], to examine cellular calcium flux in control and treated cells. Figure 7A shows that 2 hours after the start of the p tau treatment, there was a clear spike of cytoplasmic Ca^2+^. The Fluo-4 fluorescence continued to rise in the next 30 minutes whereas the untreated control showed negligible changes. Quantification of the Fluo-4 AM fluorescence from a multi-well plate confirmed the increase of Ca^2+^ (Figure 7B, bottom). Importantly, when apomorphine was present, the degree of Ca^2+^ spike was reduced (Figure 7B bar graph). Under a similar condition of early treatment, key indicators for ER stress did not change significantly (Supplemental Figure 6), which established that calcium dyshomeostasis preceded the full activation of the ER stress.

**Figure 7.**
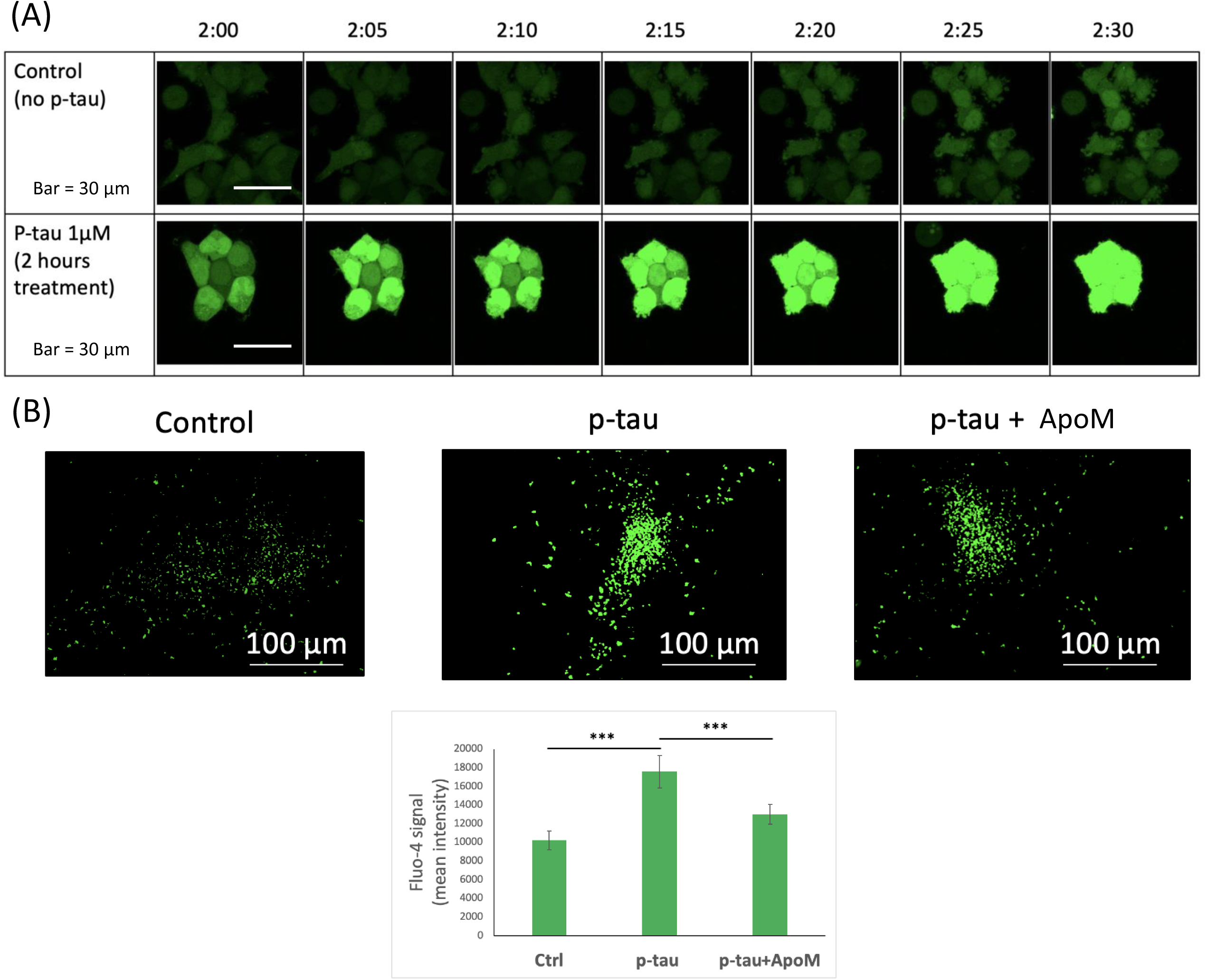
P-tau causes intracellular free calcium levels to spike that can be prevented by apomorphine. (A) SH-SY5Y cells were treated with 1 µM p-tau for 2 hours and recorded by confocal microscopy for 30 minutes. Fluo-4 was added 1 hour prior to the imaging. (B) SH-SY5Y cells were treated with 1 µM p-tau in the presence or absence of 10 µM apomorphine for 2 hours and imaged by the Cytation 5 (Biotek) cell imager at 4X magnification. Fluo-4 was added 1 hour prior to imaging. Quantification of the mean Fluo-4 intensity per cell in each sample. Error bars are SEM, n = 10. p value was determined by unpaired t test. *** P ≤ 0.001.

### Heme oxygenase-1 is a key target of p-tau attack

To gain a comprehensive view of cellular responses to p-tau, we took a non-biased proteomics approach to probe the early phase of the tau insult. Considering the wealth of proteins associated with neurofibrillary tangles[92, 93], we hypothesized that proteomic changes were part of the first wave of molecular events triggered by the cytotoxic tau. If true, identifying such changes would lead to better understanding of the disease mechanisms and possibly to promising targets for therapeutics development. To test this hypothesis, we conducted a label free, quantitative proteomic analysis by LC-MS/MS to identify proteins whose abundance was significantly altered by p-tau within 6 hours of treatment. Briefly, SH-SY5Y cells were treated with 1 µM of p-tau for 6 hours before significant morphological or viability changes were observed. Three replicas of the treated and sham control were collected, lysed with urea, and digested with trypsin before nano LC-MS/MS analysis (see Methods). Proteins quantified in all six samples were expressed in a volcano plot [-log_10_(P value) vs. log_2_(fold change)] in Figure 8. Twelve proteins increased 50% or more (red circles), one reduced 74% (HO-1), and three were down by 25% to 41% (DLD, ALDH16A1, POLA2). The top two upregulated proteins were MIOS and SYNE2 that had doubled in their abundance during the treatment.

**Figure 8.**
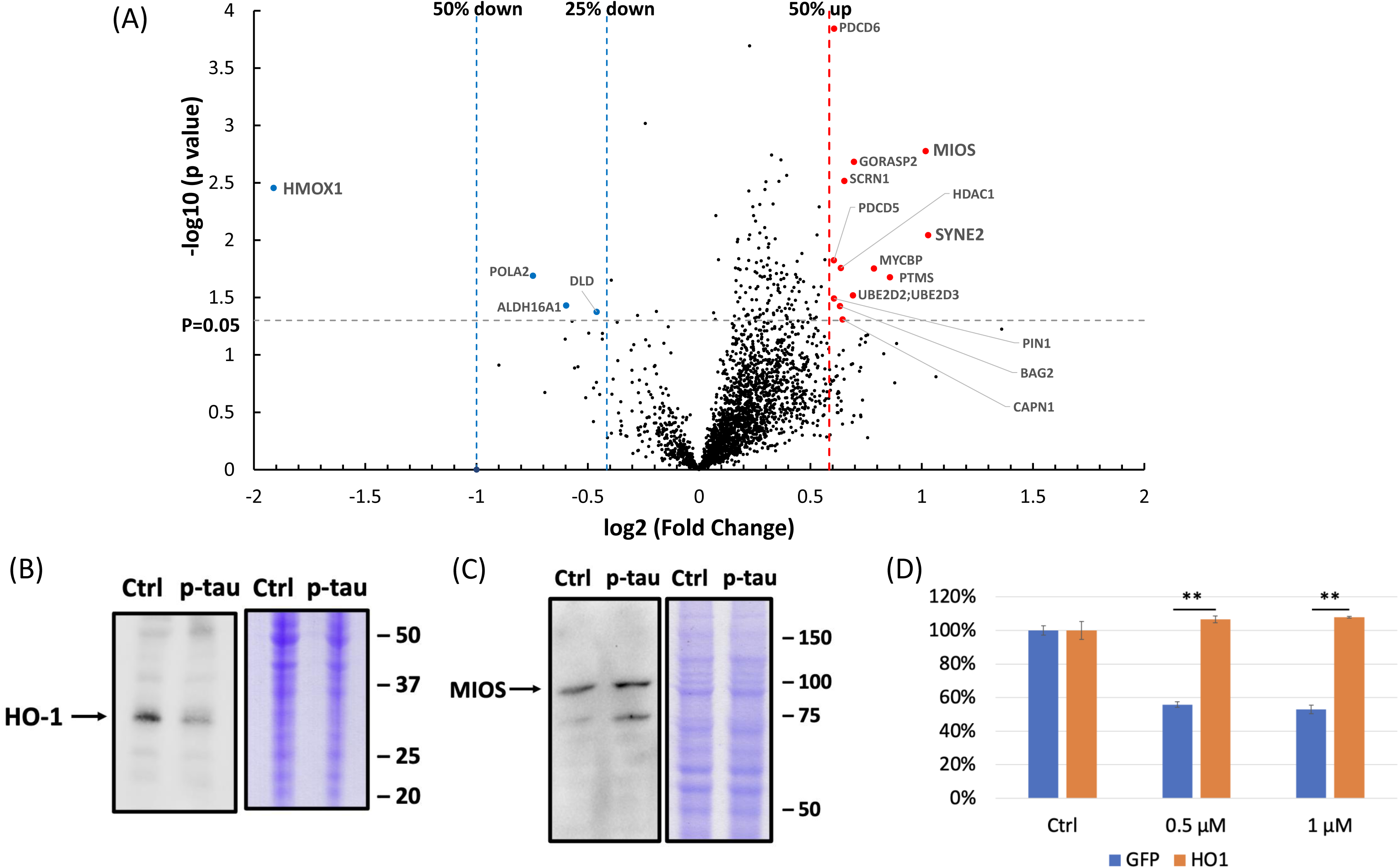
Quantitative proteomics reveals cellular molecular changes mediated by p-tau (p-tau). (A) Volcano plot shows proteomic changes in SH-SY5Y cells after 6-hour treatment of 1 µM of p tau. The x-axis is the log_2_ fold change and y-axis is the −log_10_ (P value). The vertical dot lines highlight the fold changes of -2, -1 and +2; the horizontal dot line represents a P value of 0.05; the (left-most) blue points represent the down-regulated proteins; and the (right-most) red points represent up-regulated proteins. (B) Representative immunoblots of HO-1 reduction and (C) MIOS enhancement in cell lysates after 1 µM of p-tau treatment for 6 hours (p-tau) compared with untreated control (Ctrl). Total proteins were stained with Coomassie brilliant blue as the loading control. Marker sizes are given to the right in kDa. (D) Overexpression of HO-1 confers resistance to the cytotoxicity of p-tau. The viability of SH-SY5Y cells after the overnight treatment of the indicated dose of p-tau was examined by CCK-8 assay. The numbers were normalized to each untreated group. Error bars are SEM, n = 3 biological repeats. p value was determined by unpaired t test. **p < 0.01.

The three top hits, HO-1 (heme oxygenase-1), MIOS, and SYNE2, exert diverse functions, yet each has been linked to neurodegeneration and Alzheimer’s disease[90, 94, 95]. To verify the LC-MS/MS results, immunoblotting with antibodies specific for these three proteins was done. Figures 7B and 7C confirmed the HO-1 reduction and MIOS elevation after 6 hours of treatment of p-tau. Unfortunately, we have yet to obtain conclusive results for SYNE2 due partly to the titer and specificity of the antibodies that we used.

The downregulation of heme oxygenase-1 is particularly intriguing. HO-1 is an inducible enzyme that converts the cytotoxic free heme to biliverdin in a reaction that also produces carbon monoxide (CO), Fe, and NADP [96, 97]. CO and Fe^2+^ may contribute to oxidative stress. The association between heme metabolism and neurodegeneration is well established[98, 99]. However, whether HO-1 prevents or promotes neurodegeneration is under debate[100, 101]. If HO-1 deficiency was one of the reasons underlying the cytotoxicity of p-tau, overexpressing this protein would afford protection against a cytotoxic tau. To test this hypothesis, SH-SY5Y cells were first transfected with pCX-HO1-2A-EGFP[102], from which HO-1 and eGFP were transcribed as a single chimeric mRNA. The intervening viral 2A sequence caused translation re initiation at the junction between 2A and eGFP, resulting in separate HO-1 and eGFP proteins[103]. Detection of the downstream eGFP fluorescence indicated the successful expression of HO-1. As a control, another plasmid expressing GFP but without HO-1 was used in parallel throughout the experiment. Following transfection, cells with GFP fluorescence were isolated by flow cytometry. After recovery for a day, these GFP-positive cells were treated with 0, 0.5, or 1 µM of p-tau for 20 hours before viability quantification using a colorimetric method (CCK-8). The p-tau-dependent decline of cell viability was evident in the control cells (blue bars, Figure 8D). In contrast, cells overexpressing HO-1 (orange bars) showed better survival, consistent with the hypothesis that HO-1 deficiency played a critical role in tau-mediated cell death.

In summary, data presented above revealed critical cellular functions impaired by p-tau. Elevated ER stress, unfolded protein response and inflammatory signaling pathways were likely triggers for cell death. In addition, calcium dyshomeostasis, deficiency of heme oxygenase 1, and upregulation of MIOS were among the early responses occurring before ER stress activation. Apomorphine rendered cells more resistant to the toxicity of tau. The notion that efficacious therapy can be achieved by restoring cellular functions impaired in tau pathology was supported by the observation that overexpressing HO-1 improved cell viability following p-tau attack. These data therefore provide insights into tauopathy mechanisms and suggest paths to drug development.

## Discussion

Abnormally phosphorylated tau is key to the development of Alzheimer’s disease and other neurodegenerative tauopathies[4]. Understanding the molecular basis for tau-inflicted cell dysfunction and death will facilitate the design of innovative tau-centric therapeutics. P-tau produced by the PIMAX approach possesses pathophysiologically relevant phosphorylation marks, and triggers mitochondrial superoxide accumulation and apoptosis in tissue culture cells and neurons[30]. Experimental findings presented herein further inform novel details of the molecular targets. Several stress pathways apparently were activated in response to p-tau. These included the Unfolded Protein Response, ER stress, and inflammation; all have been linked to the etiology of Alzheimer’s disease[68]. Before these detrimental events exhaust cellular viability, two early molecular responses further shed light on the action of tau. Firstly, a short, 2-hr incubation was sufficient to cause a spike of the cytosolic Ca^2+^ concentration (Figure 7). Calcium is an essential second messenger that regulates numerous cell activities in eukaryotes. Activities specific for neurons that rely on proper Ca^2+^ signaling are synaptic transmission, learning and memory formation, and long-term potentiation[104]. Dysregulation of calcium homeostasis is thought to be a critical contributor to Alzheimer’s disease[105, 106]. That p-tau causes cytosolic calcium to rise acutely suggests that calcium dyshomeostasis is an upstream stress that may lead to other damages, such as ER stress and UPR[89]. Our preliminary test on the source of this cytosolic Ca^2+^ spike is yet inconclusive, except that Dantrolene, an inhibitor of Ca^2+^ release through the ryanodine receptor (RYR) channels[107], had no effect on either Ca^2+^ spike or cell survival (data not shown), suggesting a different route of Ca^2+^ release from ER, such as the inositol-1,4,5-trisphosphate receptors (InsP_3_Rs). Alternatively, p-tau may trigger Ca^2+^ influx from the extracellular space through the voltage-gated calcium channel of the plasma membrane calcium ATPase pump[106], or a damage on the cytoplasmic membrane[31, 32]. Additional work is underway to delineate this intriguing effect of p-tau.

In addition to calcium dyshomeostasis, another early molecular consequence of p-tau treatment is the changes of the abundance of selective proteins. The most notable targets were heme oxygenase-1, MIOS and SYNE2 (Figure 8A-C). HO-1 showed 75% reduction, whereas MIOS and SYNE2 were both elevated to approximately 2-fold. HO-1 converts the pro-oxidant free heme to anti-oxidants biliverdin, CO, and Fe^2+^[99]. One attractive model explaining tau pathology is that free heme molecules resulting from a sudden restriction of HO-1 availability cause oxidative stress, which is consistent with the observation that p-tau also triggers the accumulation of mitochondrial superoxide[30]. Superoxide and other radicals then set off a cascade of stresses and eventually cell death. Critically, the hypothesis that HO-1 deficiency facilitates tau cytotoxicity is supported by the observation that overexpressing HO-1 renders cells more resistant to p-tau (Figure 8D). Taking these findings together, we propose a model in Figure 9 for cellular responses and targets for p-tau (see legend for description).

**Figure 9.**
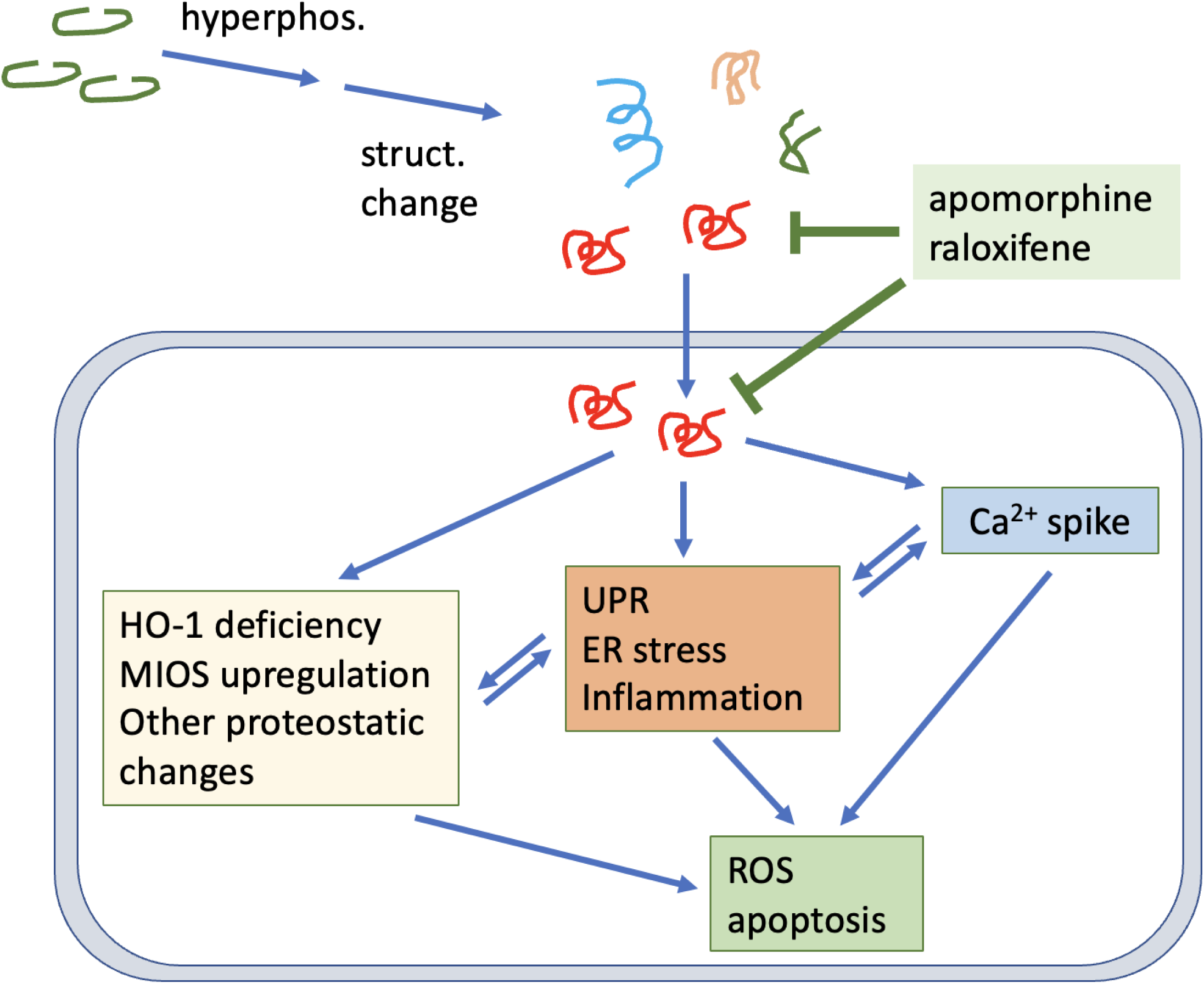
Summary illustration. P-tau triggers integrated cellular stress responses in a manner that is ameliorated by apomorphine. Tau cytotoxicity is manifested by disruption of the cytoskeleton, calcium dyshomeostasis, and the activation of Unfolded Protein Response, ER stress and inflammation pathways. In addition, proteomic changes immediately following the insult of p-tau include the upregulation of MIOS in the mTOR pathway for energy balance, and the downregulation of HO-1 essential to keep heme at bay. Cells may die of one or more molecular defects, including oxidative damages and apoptosis. Apomorphine and HO-1 are labelled red for their apparent efficacy of protecting cells from p-tau cytotoxicity. Apomorphine likely exerts its anti-tau pathology function by directly modulating the conformation of tau, as such, cellular pathways instigated by p-tau are all effectively restored by this compound. Easing the HO-1 deficiency by overexpressing this protein results in enhanced resistance to p-tau. These two ameliorative approaches suggests that efficacious treatment of Alzheimer’s disease may be achieved by combatting the neurotoxic tau, and by galvanizing cellular functions crippled by the pathological tau.

Arguably, one of the most impactful outcomes from tau-centric AD research will be the development of novel therapeutics that prevents or slows down the neurodegeneration caused by abnormally phosphorylated tau. Given that p-tau wreaks havoc on cultured cells in a timeframe seemingly too short to form the canonical fibrils that constitute neurofibrillary tangles (Figure 2, and see[30]), and that tangles and cognitive decline are in large separable in animal models (e.g.,[29, 108, 109]), we argue that measures curbing the cytotoxicity, instead of fibrillogenesis, of tau is a more viable approach to drug development. For example, methylene blue and its derivatives[110] were effective in preventing paired helical filament formation and dissolving pre-formed tangles in vitro, but failed to improve cognition in clinical trials[55, 56], a disappointing outcome reminiscent of the inability of Aβ-targeting drugs to robustly slow down the cognitive decline of patients[111], despite that the Aβ burden was significantly reduced. A high-throughput screen for tau aggregation inhibitors using the K18 fragment of unmodified tau identified primarily redox modulators with limited therapeutic potential[47, 53], suggesting that the canonical aggregation assays of unmodified tau may miss a key pathological element(s). On the contrary, apomorphine was identified in a pilot screen for compounds that antagonized both amyloid genesis and cytotoxicity of p-tau[58]. We showed here that not only did apomorphine control the higher-ordered structure of the pre-tangle tau (Figure 2) and the overall amyloid content (Supplemental Figure 5), but it also stopped essentially all the tau instigated stress responses tested in this work. Intriguingly, internalization of p-tau appeared to be unaffected by apomorphine (Figure 3B), consistent with our previous report that treating cells with apomorphine 24 hours after the addition of p-tau was still effective in reducing cell death[58]. These observations suggest a window of opportunity of treatment between the uptake of pathological tau and the death of the underlying cells.

The non-biased proteomics analysis for proteins that responded to p-tau insult within six hours uncovered intriguing candidates. Of those that showed the most significant enrichment included MIOS and SYNE2. MIOS is a component of the conserved GATOR2 (a.k.a. SEACAT) complex that activates the mammalian target of rapamycin (mTOR) pathway when amino acid nutrients are present[112]. mTOR is a major intracellular signaling hub that responds to nutrients, growth factors and mitogens. The simplest model for our proteomics finding is that the p-tau-induced upregulation of MIOS promotes the action of GATOR2, which in turn activates mTOR. P-tau may therefore impact the energy and metabolism balance of cells. Inhibiting mTOR by rapamycin results in the extension of lifespan and healthspan of experimental animals[113]. Thanks to this anti-aging activity, rapamycin is currently tested in a phase 2 clinical trial (REACH, https://clinicaltrials.gov/ct2/show/study/NCT04629495) for the therapeutic potential against mild cognitive impairments, for aging is the most important risk factor for AD. It will be interesting to see whether rapamycin exerts an inhibitory activity against p-tau in vitro. In addition, the mTOR pathway appears to be a pivot that links cellular energetics and metabolism to both Alzheimer’s disease and type 2 diabetes (T2D)[94, 114]. T2D patients have a 2-fold higher risks of getting AD[115]. The upregulation of MIOS may provide a causal link to this path of neuron loss. SYNE2 encodes the Nesprin protein 2 that is associated with laminopathies in human, most notably the Emery-Dreifuss muscular dystrophy[116]. Additionally, Nesprins are implicated in other diseases including cancer, hearing loss and neurological disorders[117]. The Nesprins complex coordinates the physical and functional association between the nuclear membrane and cytoskeleton[117, 118]. Whether the cytoskeleton disorganization seen in Figure 6 was connected to SYNE2 is an interesting topic for further investigation.

Heme oxygenase-1 suffered the most severe deficit within six hours of the treatment with p-tau, hinting heme metabolism as a key immediate response to the tau insult. HO-1 is an inducible enzyme that converts the pro-oxidant free heme molecules to anti-oxidants biliverdin and bilirubin[101]. One possible underpinning of tau pathology is therefore an imbalanced redux microenvironment in cells. Although we have yet to obtain convincing evidence for significant accumulation of heme in cells attacked by p-tau, that overexpressing HO-1 empowered the underlying cells to resist tau cytotoxicity (Figure 8) lends strong support to targeting the cellular functions impaired by pathological tau as a therapeutic goal. Given the involvement of ER stress and UPR, measures that restore proteostasis through, for example, regulating the activity of certain heat shock proteins appear to be a worthy direction for AD drug development[119].

In addition to selective pathways impaired by pathological tau, targeting the root of the pathology, i.e., p-tau, with apomorphine and possibly other small-molecule compounds affords an attractive option for AD intervention. Apomorphine controls the three-dimensional structure of p-tau (Figure 2) and reduced the cytotoxicity of p-tau (this work and [58]). Detailed mechanistic studies shown in this work showcase the powerful therapeutic potential of apomorphine. All molecular defects inflicted by p-tau, including ER stress, UPR, inflammation, calcium dyshomeostasis, and cytoskeleton disintegration, were effectively suppressed by apomorphine. In the screen that identified apomorphine as an effective inhibitor for the aggregation and cytotoxicity of p-tau, we also identified raloxifene with comparable protective power as that of apomorphine[58]. Indeed, raloxifene also modulates the high-ordered structure of p-tau (see Supplemental Figure 4). Whereas we did not test whether raloxifene had a similar spectrum of cytoprotective effects, there is no reason to speculate the otherwise. Both apomorphine and raloxifene are prescription drugs with well-characterized pharmacological characters[120, 121]. Raloxifene reduced the risks of mild cognitive impairments by 33% in a 5-year, randomized, placebo-controlled trial of 5,386 postmenopausal women with osteoporosis[61]. However, in a subsequent 1-year trial on older women already showing mild to moderate cognitive symptoms, this compound was deemed ineffective in preserving cognition[122]. Apomorphine was shown to improve the memory of 3xTg AD mice that express both Aβ and tau[60] by enhancing the insulin signaling pathway and reducing Aβ burden. However, it remains formally possible that apomorphine injected to 3xTg mice antagonized the pathological activity of the transgenic tau. The improved insulin signaling pathway may actually result from the subdued tau toxicity. Given the long duration and gradual progression of AD pathology before cognitive symptoms arise, it is very difficult to pinpoint the ideal time to start the practical treatment, if a drug is available. In contrast to AD, traumatic brain injury, a tauopathy with high risks of developing dementia[123], has a definitive time of onset. Therefore, before a reliable early AD biomarker that informs the time of early treatment, apomorphine and raloxifene seem to be excellent candidates for clinical trials on TBI patients to stop the progression of damages caused by abnormally phosphorylated tau following the impact.

Alzheimer’s disease is a complicated neurological disorder involving many cell types and brain areas. The interplays of different tissues and organs are beginning to be appreciated by researchers. In vitro studies shown in this work afford a fundamental view of molecular and cellular alterations triggered by p-tau. Despite that all cells that we have tested so far exhibited significant and likely overlapping responses to p-tau, it is very likely that cell-specific functions may be differentially affected by p-tau. Three-dimensional cultures, organoids, and new animal models will provide the platforms for thorough pathophysiological research of Alzheimer’s disease and other tauopathies.

## Conclusion

P-tau is a potent trigger of several stress responses such as the Unfolded Protein Response, ER stress, and inflammation that have been linked to Alzheimer’s disease. Prior to the full activation of the above stress responses and cell death, a spike of the cytosolic calcium and the deficiency of heme oxygenase-1 are observed. A small-molecule compound, apomorphine, that controls the high-ordered structure of p-tau also prevents the above stress responses. Consistent with the proteomics findings, overexpressing heme oxygenase-1 confers cellular resistance to p-tau. Together, these findings provide novel and efficacious means of treating Alzheimer’s disease.

## List of Abbreviation

Aβ: amyloid β
AD: Alzehiemer’s disease
ApoM: apomorphine
ATF: activating transcription factor
CAVI: carbonic anhydrase VI
CCK-8: cell counting kit-8
CHOP: C/EBP homologous protein
CO: carbon monoxide
DR5: death receptor 5
ER: endoplasmic reticulum
GATOR: GAP activity toward RAGs (GATOR) complex
GFP: green fluorescence protein
GSK-3β: glycogen synthase kinase-3β
HO-1: heme oxygenase-1
InsP_3_Rs: inositol-1,4,5-trisphosphate receptors
IRE1α: inositol-requiring enzyme 1α
JNK: JUN N-terminal kinase
LC-MS/MS: liquid chromatography - tandem mass spectrometry
MIOS: missing oocyte, meiosis regulator, homolog
mTOR: mammalian Target of Rapamycin
NADP: oxidized nicotinamide adenine dinucleotide phosphate
NFκB: nuclear factor-κB
NFT: neurofibrillary tangle
PERK: protein kinase-like endoplasmic reticulum kinase
PHF: paired helical filament
ROS: reactive oxygen species
SEACAT: activation of Seh1-associated complex
SYNE2: synaptic nuclear envelope protein 2
T2D: type 2 diabetes
TRB3: Tribbles homolog 3
TEM: transmission electron microscopy
UPR: Unfolded Protein Response

## Declarations

### Ethics approval and consent to participate

Not applicable

### Consent for publication

Not applicable

### Availability of data and materials

The datasets used and/or analyzed during the current study are available from the two corresponding authors on reasonable request.

## Authors’ contributions

HTCH and HRC were responsible for data shown in Figure 1. ZS, K-WW, and HTCH were responsible for data shown in Figures 2C, 4 and 5 (ZS), Figures 2A, 2B, 3 and 6 (KWW), and Figure 7 (H-TCH). CYK, KZ and MHK oversaw the overall progression of the research. All authors participated in the writing of the paper.

## Acknowledgements

We thank Stacy Hovde, Sandhya Payankaulum, Nicholas Lewis, Mengyu Liu, Hsin-ying Lin, and Kai-Ching Hsiao for discussion and comments throughout the progression of this work. This work was supported by NIH grants (AG062435, AG057274) to MHK, and NIH grants (DK090313, DK126908, and DK132065) to KZ.

---

**Supplemental Figure 1.**
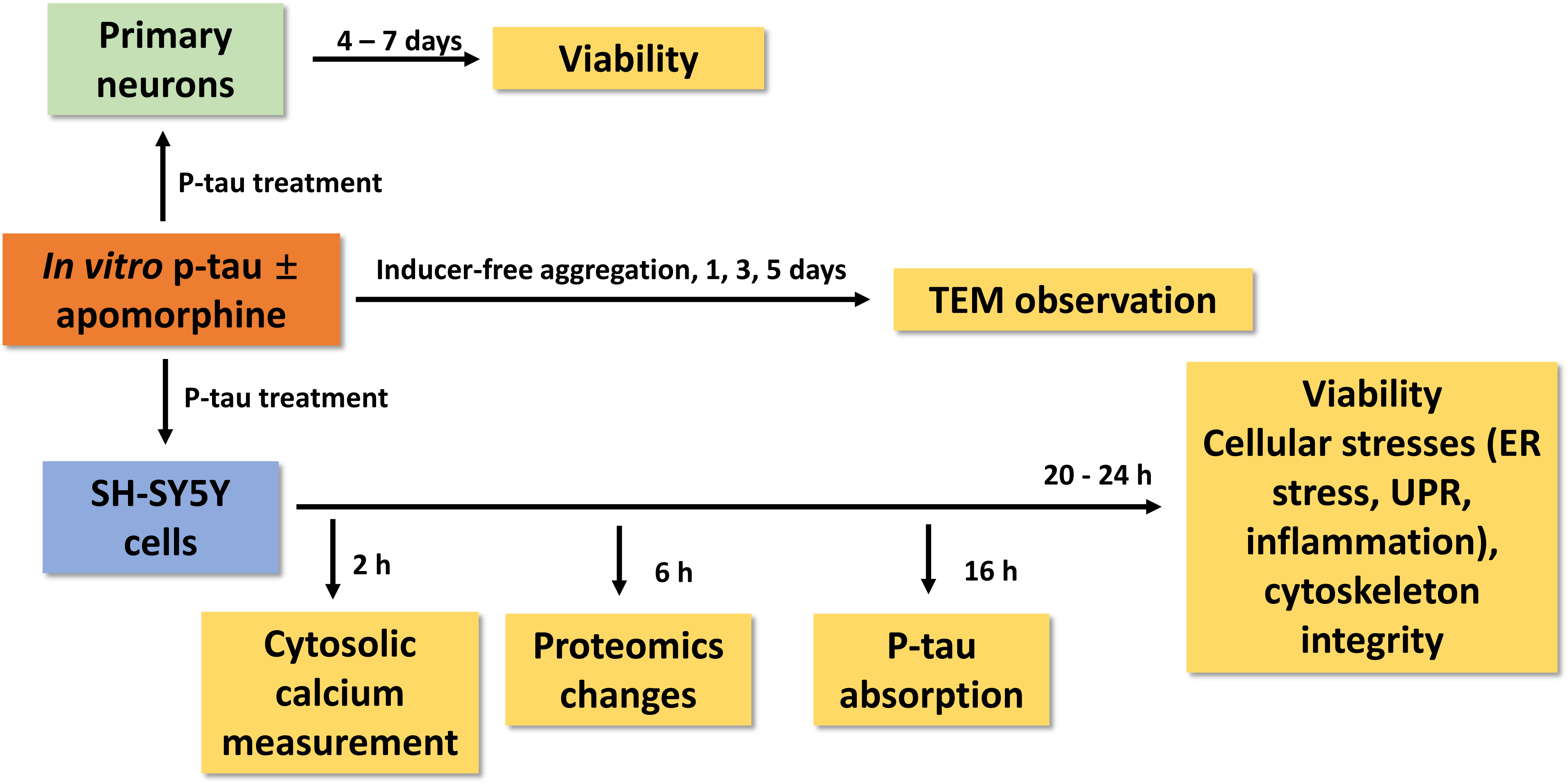
Overall design and time course of experiments shown in this work.

**Supplemental Figure 2.**
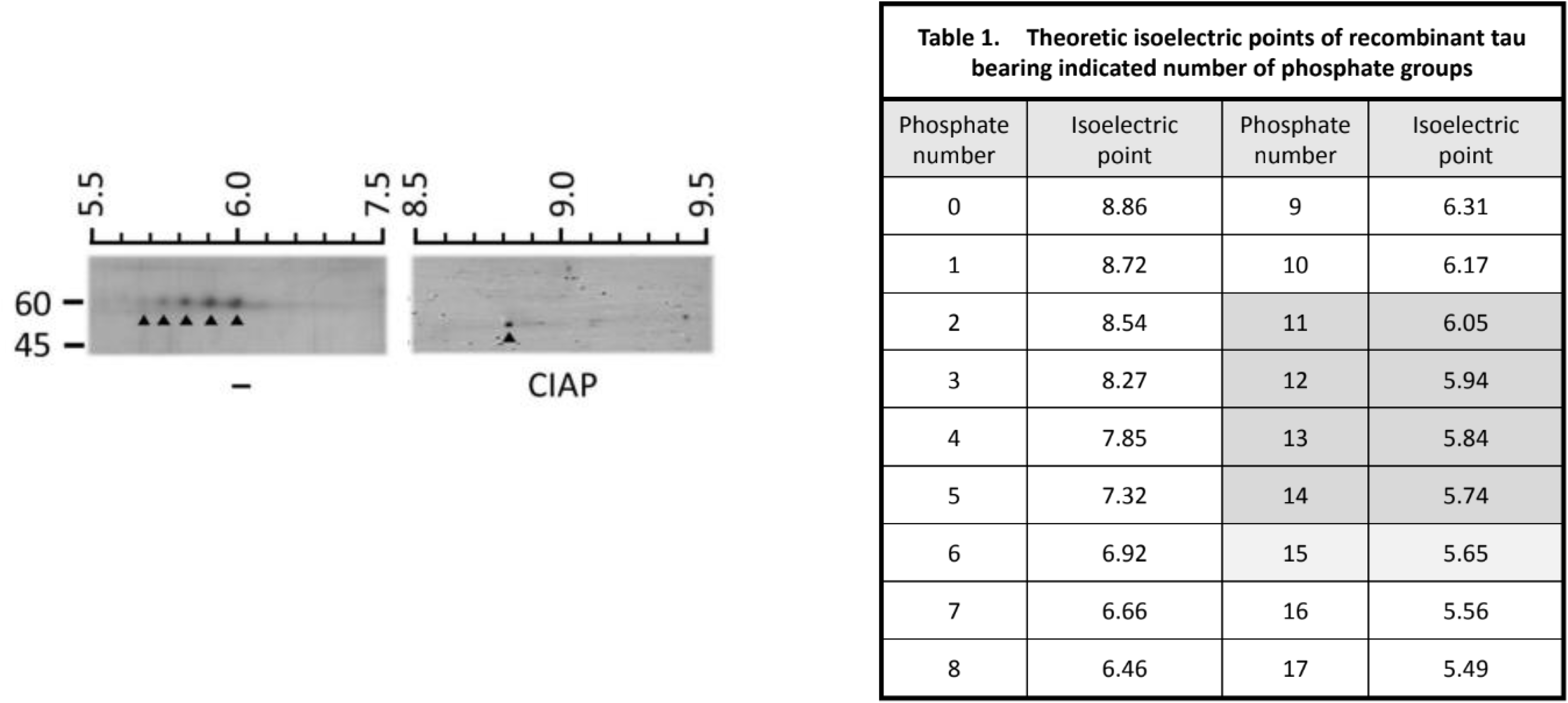
Isoelectric focusing-SDS-PAGE two-dimensional gel electrophoreses of p-tau with (right) or without (left) calf intestinal alkaline phosphatase treatment (0.5 U, 37°C, 0.5 hr). P-tau was resolved with IPG strip with pH 4-7 (left) or pH 6-11 (right) gradient, followed by SDS-PAGE (10%) that resolved proteins by size. The number to the left showed the migrated positions of MW standard proteins. The numbers at the top are the putative pH at indicated positions of IPG gel strips. Upward arrow heads mark the positions of different phosphorylated tau species.

**Supplemental Figure 3.**
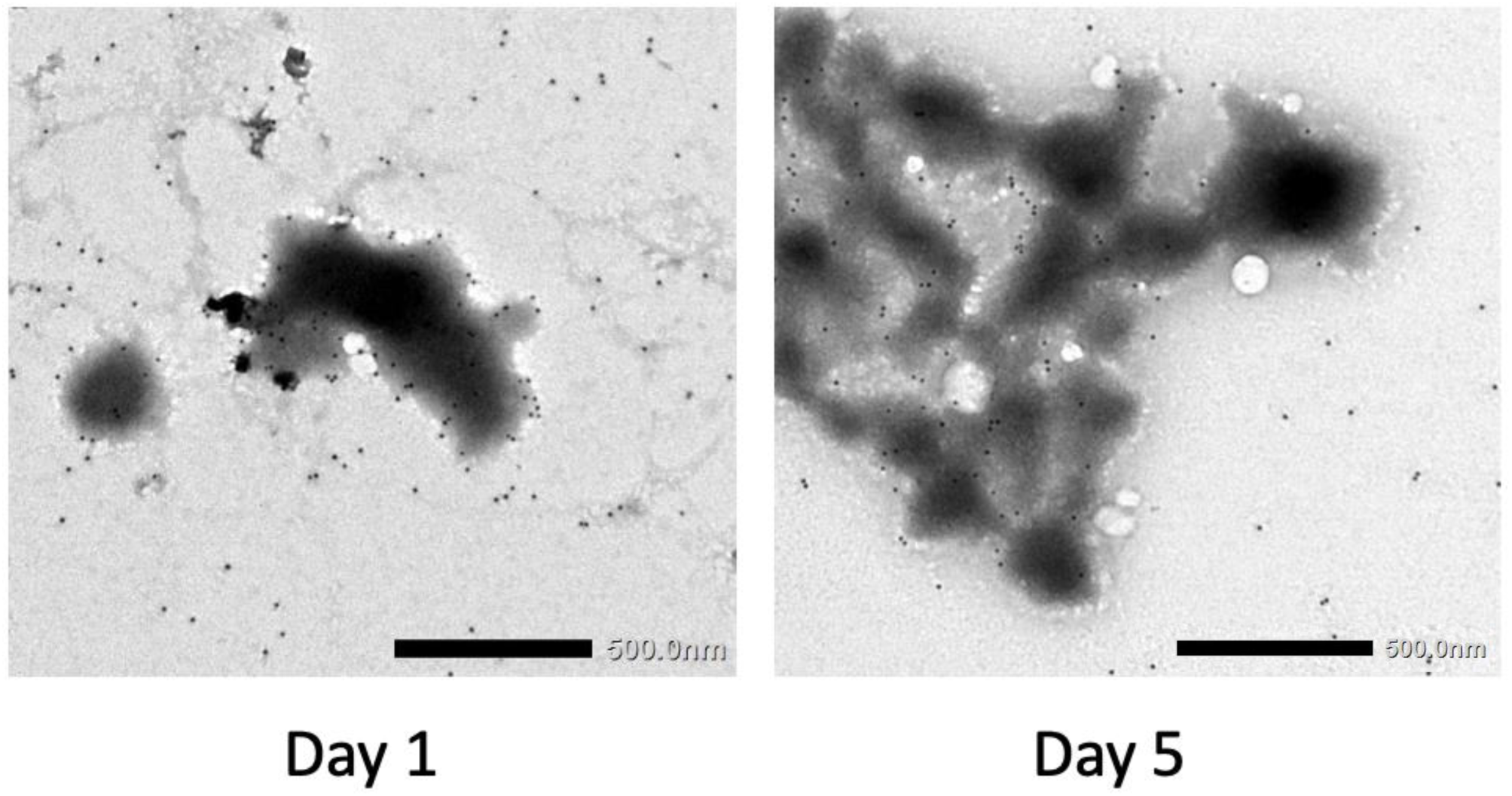
The amorphous aggregates are composed of tau. 1 µM p-tau (p-tau) was fix on the grid and recognized by DA9 tau antibody followed by immunogold particles. The sample was then negatively stained by uranyl acetate and imaged by TEM. Scale bar = 500 nm. Note that the gold nanoparticles tend to stain the outer edge of large structure, suggesting that the core of these large complexes might be very dense so that the epitope was masked. In contrast, smaller, structures with lower electron density were easily stained, suggesting better accessibility.

**Supplemental Figure 4.**
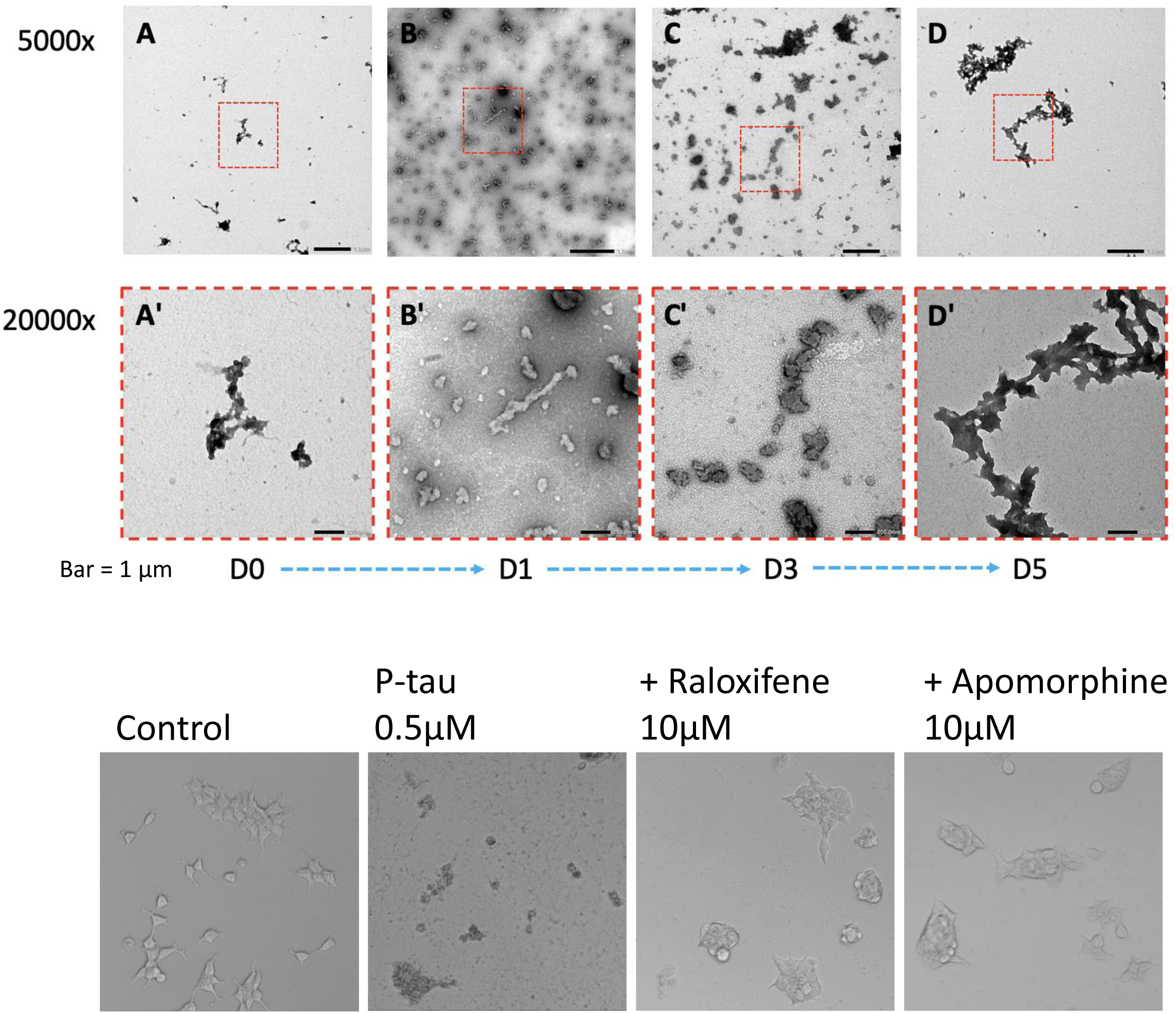
Raloxifene also controls the higher-ordered structure of p-tau in vitro. The effects of raloxifene during a 5-day aggregation reaction of p-tau were examined along with apomorphine (see Figure 1 legend for details). While both compounds altered the structures of tau aggregates, there appeared to be differences in the overall size and morphology, as shown by high-magnification images.

**Supplemental Figure 5.**
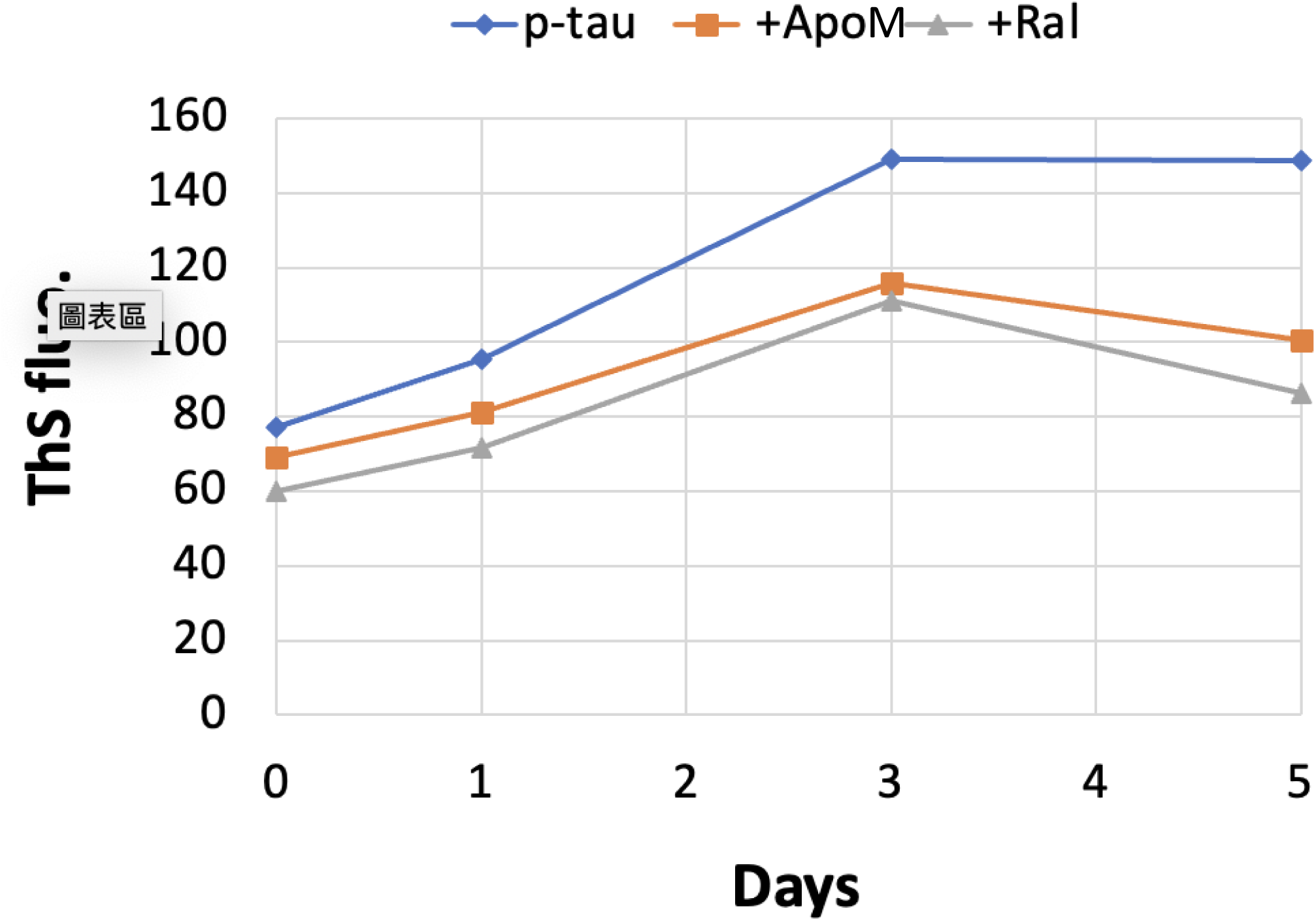
The amyloid content of p-tau (p-tau) continued to increase for up to five days, while apomorphine and raloxifene maintained their inhibition potency. 10 µM p-tau aggregation assay was done at 37 degrees in the absence of thioflavin S (ThS). At the indicated time, a portion of the reaction was removed and mixed with 20 µM ThS before fluorescence measurement.

**Supplemental Figure 6.**
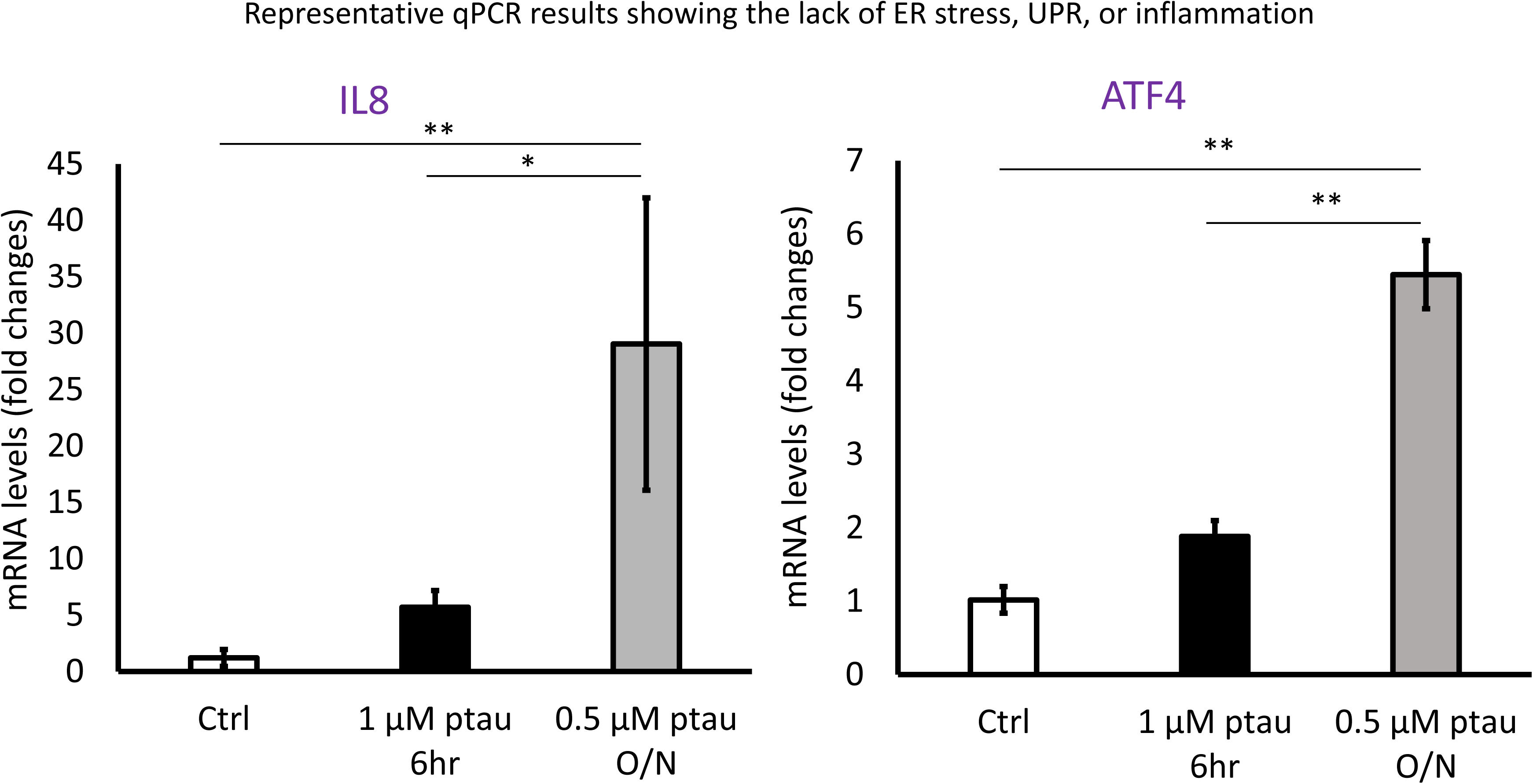
Short-term incubation with p-tau (p-tau) doesn’t induce inflammation and ER stress. IL8 and ATF4, representative inflammation and ER stress indicators, were activated in long-term p-tau treatment while not in the short-term course. Expression of the indicated ER stress-related genes was determined by qRT-PCR from SH-SY5Y cells with 6-hour 1 µM p-tau or overnight 0.5 µM p-tau treatment. The numbers were normalized to GAPDH expression. Error bars are standard deviation of biological repeats, n = 3. p value was sdetermined by unpaired t test. ** P ≤ 0.01.

