## Supplemental table 1 for "Hyperphosphorylated tau Inflicts Intracellular Stress Responses That Are Mitigated by Apomorphine"

**S-Table 1.** Primer sequence information for qPCR used in this study

| **Gene name** | **Forward primer** | **Reverse primer** |
| --- | --- | --- |
| IRE1α | ATT GTG TAC CGG GGC ATG TT | CTC ACG GTC TGC GAA GCT AA |
| XBP1s | CCGCAGCAGGTGCAGG | GAGTCAATACCGCCAGAATCCA |
| P58ipk | GGTGCTGAATGTGGAGTAAATGC | AGTAGCCCTCCGATAATAAGCAA |
| ERO1b | TTCTGGATGATTGCTTGTGTGAT | GGTCGCTTCAGATTAACCTTGT |
| ATF3 | GAGGATTTTGCTAACCTGACGC | CTACCTCGGCTTTTGTGATGG |
| ATF4 | TCAAACCTCATGGGTTCTCCA | CACAGCCAGCCATTCGG |
| DR5 | GCCCCACAACAAAAGAGGTC | AGGTCATTCCAGTGAGTGCTA |
| TRB3 | TACCTGCAAGGTGTACCCC | GGTCCGAGTGAAAAAGGCGTA |
| MCP1 | TGCAATCAATGCCCCAGTCA | GCTTCTTTGGGACACTTGCTG |
| TNFa | CCCATCTATCTGGGAGGGGT | ATCCCAAAGTAGACCTGCCC |
| IL8 | TCTGCAGCTCTGTGTGAAGG | TGGGGTGGAAAGGTTTGGAG |
| INFγ | CCAAGTGATGGCTGAACTGTC | TACTGGGATGCTCTTCGACCT |
| GADD34 | ATGATGGCATGTATGGTGAGC | AACCTTGCAGTGTCCTTATCAG |
| GAPDH | TGAAGGTCGGAGTCAACGG | AGAGTTAAAAGCAGCCCTGGTG |
